# An evolutionarily conserved laterally acquired toolkit enables microbiota targeting by *Trichomonas*

**DOI:** 10.1101/2025.04.21.649813

**Authors:** Adam J. Hart, Lenshina A. Mpeyako, Nick P. Bailey, George Merces, Joseph Gray, Jacob Biboy, Manuel Banzhaf, Waldemar Vollmer, Robert P. Hirt

## Abstract

*Trichomonas* species are a diverse group of obligate extracellular symbionts associated with or attributed to various inflammatory diseases. They colonise mucosal surfaces across a wide range of hosts, all of which harbour a resident microbiota. Their evolutionary history likely involved multiple host transfers, including zoonotic events from columbiform birds to mammals. Using comparative transcriptomics, this study examines *Trichomonas gallinae* co-cultured with *Escherichia coli*, identifying a molecular toolkit that *Trichomonas* species may use to interact with bacterial members of the microbiota. Integrating transcriptomic data with comparative genomics and phylogenetics revealed a conserved repertoire of protein-coding genes likely acquired through multiple lateral gene transfers (LGT) in a columbiform-infecting ancestor. These LGT-derived genes encode muramidases, glucosaminidases, and antimicrobial peptides—enzymes and effectors capable of targeting bacterial cell walls, potentially affecting the bacterial microbiota composition across both avian and mammalian hosts. This molecular toolkit suggests that *Trichomonas* species can actively compete with and exploit their surrounding microbiota for nutrients, potentially contributing to the dysbiosis associated with *Trichomonas* infections. Their ability to target bacterial populations at mucosal surfaces provides insight into how *Trichomonas* species may have adapted to diverse hosts and how they could influence inflammatory mucosal diseases in birds and mammals.

## Introduction

*Trichomonas spp.*, are obligate extracellular symbionts of animal mucosal surfaces where they exhibit specific interactions with their hosts and the members of the resident microbiota (Maritz *et al*., 2014). *Trichomonas vaginalis* and *T. gallinae* are also known to harbour microbiotas of their own, sheltering some mycoplasma-like species (Margarita *et al*., 2020; Bailey *et al*., 2023) and some RNA viruses (Graves *et al*., 2019). The interplay between these symbiotic relationships is thought to impact pathogenesis and disease (Mercer & Johnson, 2018; Hirt, 2019; Margarita *et al*., 2020). Infections with *Trichomonas* species are often asymptomatic (Ribeiro *et al*., 2015; Qiu *et al*., 2017; World Health Organisation, 2023), and when disease is observed it is typically accompanied by evidence of dysbiosis (Brotman *et al*., 2012; Bisson *et al*., 2019; Ji *et al*., 2020). Microbiota dysbiosis is a disease-promoting functional state of a site’s microbial population that contrasts with an optimal homeostasis-promoting, eubiotic, microbial population (DeGruttola *et al*., 2016; Hooks & O’Malley, 2017; Levy *et al*., 2017; Hou *et al*., 2022). Dysbiosis is typically associated with increased inflammation and making the host prone to further infections (DeGruttola *et al*., 2016). The cause of the shift from asymptomatic to symptomatic *Trichomonas* species infection and dysbiosis is unclear. However, interactions between *Trichomonas* species with host cells and the microbiota are likely important factors (Hirt, 2013; Bär *et al*., 2015; Mercer & Johnson, 2018).

*T. vaginalis*, the best studied *Trichomonas* species, has several aspects of its interactions with host cells, microbiota and endosymbionts characterised (Mercer & Johnson, 2018; Margarita *et al*., 2020; Riestra *et al*., 2022). *T. vaginalis* is associated with a type IV community state type (CST) vaginal microbiota, marked by increased bacterial diversity, including increased density of strict anaerobes like *Prevotella* species, mycoplasma-like species and reduced density of mutualist *Lactobacillus* species (Ravel *et al*., 2011; Brotman *et al*., 2012; Fettweis *et al*., 2014; Saraf *et al*., 2021). Studies investigating the prevalence of *T. vaginalis* in females frequently report that infected individuals also present with co-pathogens and/or bacterial vaginosis (Krashin *et al*., 2010; Brotman *et al*., 2012; Javanbakht *et al*., 2013; Fettweis *et al*., 2014; Mitchell *et al*., 2014; de Waaij *et al*., 2017; Tchankoni *et al*., 2021; Zhu *et al*., 2023). It is suggested that the combined action of *T. vaginalis* and the dysbiotic bacterial population amplifies the negative health implications for the host organism, including combined negative immune modulation and epithelial cell permeability (Fichorova *et al*., 2013; Hinderfeld *et al*., 2019; Riestra *et al*., 2022). Notably, infections of *T. gallinae* also occur alongside the depletion of *Lactobacillus* species in pigeons’ upper digestive tracts and an increase in bacterial diversity, including anaerobes like *Prevotella* and mycoplasma-like species, but a decrease in overall bacterial abundance in the same sites (Ji *et al*., 2020; Bailey *et al*., 2023). Similarly, *Trichomonas tenax* is associated with periodontitis - an inflammatory disease driven by bacterial pathobionts in the dysbiotic environment of periodontal pockets - and is thought to contribute to this dysbiosis (Bisson *et al*., 2019). Hence, all current *Trichomonas* species infections are associated with dysbiosis in both birds and mammals.

The presence of mutualistic *Lactobacillus* species within the human vaginal mucosal microbiota is considered to have important beneficial protective capabilities via microbial competition, the lowering of vaginal pH (production of lactic acid), producing antimicrobial substances such as anti-microbial peptides (AMPs) and the regulation of the host immune system (Valenti *et al*., 2018; Han *et al*., 2021). Notably, *T. vaginalis* is associated with the depletion of *Lactobacillus* species, which the parasite could be specifically targeting to ensure the environment is optimal(Fichorova *et al*., 2013). Alternatively, pre-existing bacterial vaginosis may also create a more favourable environment for *T. vaginalis* infections. Therefore, the question remains as to whether *T. vaginalis* infection drives dysbiosis or dysbiosis promotes *Trichomonas* (Margarita *et al*., 2020).

The species members of the *Trichomonas* genus and related Trichomonads have experienced zoonotic (and possibly anthroponotic) events, posing potential global health risks (Maritz *et al*., 2014; Hirt & Sherrard, 2015). The molecular and cellular mechanisms facilitating such host transfers are not well understood. Human infecting *Trichomonas* likely originated from members of the Columbidae family (columbine birds, pigeons and doves) where known *Trichomonas* molecular diversity has been dramatically expanded by investigating Australasian columbines species, further supporting multiple host transfers, including zoonoses, from distinct birds sources (Peters *et al*., 2020). Hence, investigations into the biology of the pigeon infecting *T. gallinae* could help elucidate the mechanisms behind zoonotic transmissions of *Trichomonas* species (Maritz *et al*., 2014).

Over one hundred candidate genes originating from LGT in *Trichomonas spp.* were first identified within the genome of *T. vaginalis* (Carlton *et al*., 2007; Alsmark *et al*., 2013). While LGTs are very commonly observed among prokaryotes they are also increasingly identified among eukaryotes (Metcalf *et al*., 2014; Husnik & McCutcheon, 2018). Indeed, it appears that protists, including parasites, are some of the eukaryotes most affected by LGT (Hirt *et al*., 2015; Husnik & McCutcheon, 2018; Sibbald *et al*., 2020). Conserved genes originating from LGT can confer a significant selective fitness advantage to the recipient organism (Sibbald *et al*., 2020). Indeed, a group of laterally acquired genes encoding peptidoglycan (PG) hydrolases (NlpC/P60 peptidases), initially identified in *T. vaginalis*, and subsequently demonstrated to be shared with *T. gallinae* and *T. tenax*, were shown to be of functional importance for *T. vaginalis-*bacteria interactions (Pinheiro *et al*., 2018; Barnett *et al*., 2023). These enzymes hydrolyse PG cross-links and their transcripts are significantly upregulated in the presence of *Lactobacillus gasseri* or *E. coli* (Pinheiro *et al*., 2018; Barnett *et al*., 2023). These hydrolases however were not sufficient to lyse *E. coli* on their own in an *in vitro* assay, implying that they likely require functional partners, such as lysozymes and AMPs, to effectively access and target the members of the vaginal microbiota (Pinheiro *et al*., 2018; Barnett *et al*., 2023).

Here, we integrated cell biology, phylogenetics, and comparative transcriptomics and genomics to investigate the evolution and the cellular and molecular mechanisms underlying *Trichomonas*-bacteria interactions. This approach led to the identification of a conserved molecular toolkit - encoded by genes likely acquired through multiple LGT events from bacterial donors - that targets bacterial PG and its precursor. These proteins may contribute to the promotion or exacerbation of bacterial dysbiosis, facilitate host switching (including zoonotic transmission), and play a role in inflammatory tissue damage.

## Results

### The presence of *E. coli* increases the fitness of *Trichomonas gallinae in vitro*

To assess the possible fitness effect of bacteria on *T. gallinae*, *E. coli* (DH5α) and *T. gallinae* (strain A1, isolate XT-1081/07, clone GF1c) (Alrefaei *et al*., 2019) were co-cultured in TYM media for 48 hours at an initial 10:1 parasite-to-bacteria ratio. Hemocytometry showed that the *T. gallinae* density was significantly higher (p-value < 0.05) in monoculture than in co-culture after 24 hours (**Figure 1A**). However, this trend reversed by 48 hours, with *T. gallinae* counts significantly exceeding (≈2.3-fold) those in monoculture (p-value < 0.05). The presence of the stationary-phase or death-phase bacteria (**Figure S1**) could therefore contribute to the higher cell density of *T. gallinae* in this system.

**Figure 1.**
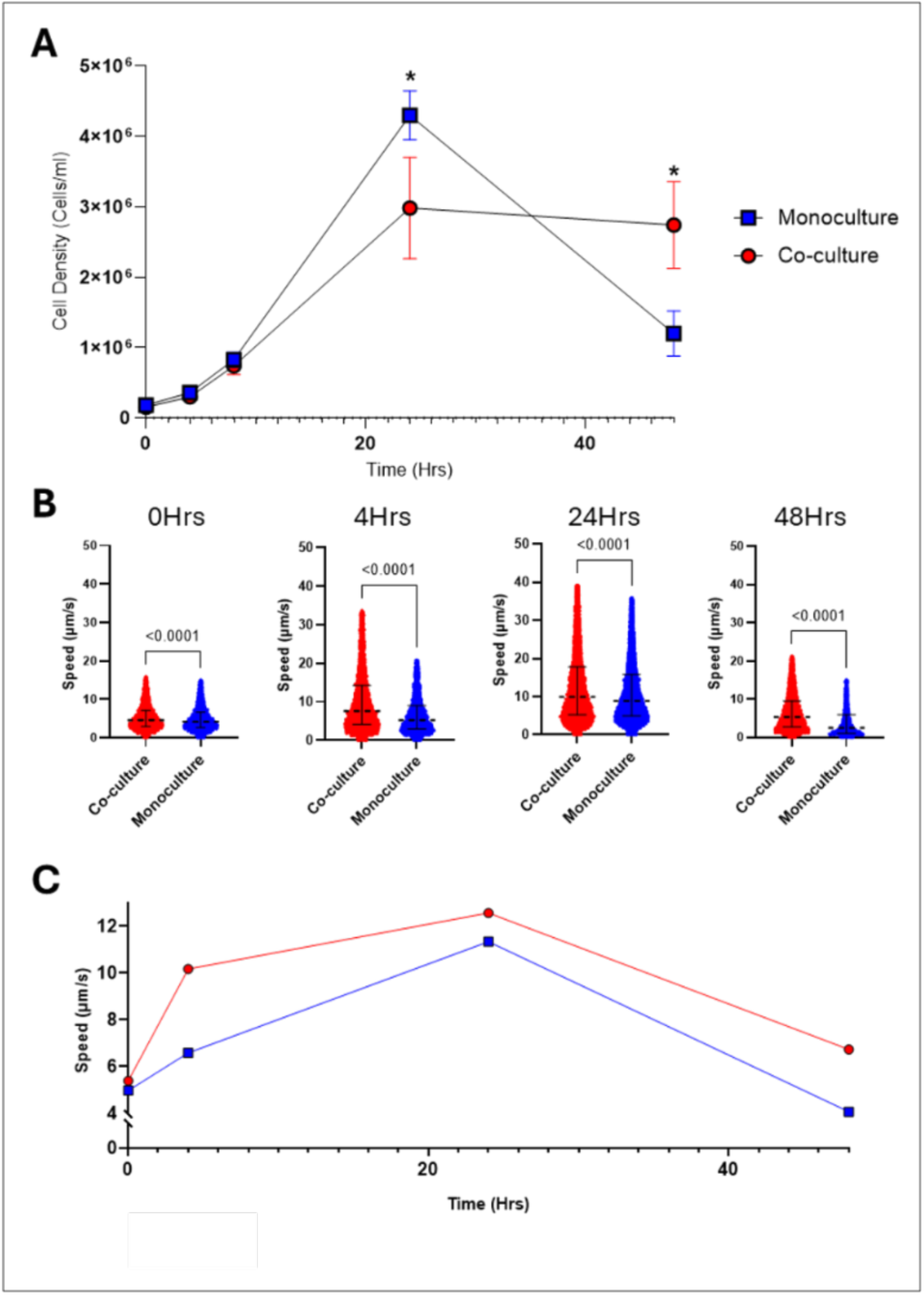
Growth and motility of *T. gallinae* in co-culture with *E. coli.* (A) Growth curve of *T. gallinae* (cells/ml) over 48 hours in monoculture (blue) and co-culture with *E. coli* (red)(n=3, error bars are standard deviations). Asterisks indicate significant differences (p-values < 0.05) between conditions (unpaired *t*-test). (B) Violin plots showing *T. gallinae* speed (µm/s) at multiple time points, measured via live-cell imaging in the absence (blue) or presence (red) of *E. coli*. P-values from Kruskal–Wallis tests are shown above each comparison. (C) Mean *T. gallinae* speed (µm/s) over time in monoculture (blue) and co-culture (red) conditions.

To further assess the fitness changes of *T. gallinae* when grown alongside *E. coli* we designed a novel cell tracking methodology/pipeline to specifically quantify *T. gallinae* motility - live cell imaging described in the Materials and Methods section. The output metrics include track duration (frames), mean speed (µm/s), total distance travelled (µm), and maximum distance travelled (µm).

This pipeline selectively tracks *T. gallinae* cells over *E. coli* in co-culture samples (**Figure S2**). *T. gallinae* in co-culture moved significantly (p-value < 0.05) faster than in monoculture across all timepoints (**Figure 1B**), with, for example, a median speed of 10.2 µm/s versus 6.6 µm/s after 4 hours (**Figure 1C**). The total distance travelled results corroborate the increase in speed (**Figure S3**). The presence of *E. coli* thus enhances *T. gallinae* motility and fitness *in vitro*.

### *E. coli* cells are partially engulfed by *Trichomonas gallinae* during ‘Pocket’ physical interactions

*Trichomonas spp.* are known to phagocytose certain bacteria and fungi; thus, transmission electron microscopy (TEM) was used to examine *T. gallinae*–*E. coli* co-cultures for phagocytosis. While no phagocytosis was observed, there was evidence for the two species interacting in close proximity, with *T. gallinae* forming invaginations containing *E. coli* cells after 4 hours of co-incubation (**Figure 2**). These invaginations, which we termed ‘pocket interactions,’ form tight localised interfaces between the *E. coli* cell envelope and *T. gallinae* membrane, representing a likely site for specific cell-cell interactions.

**Figure 2.**
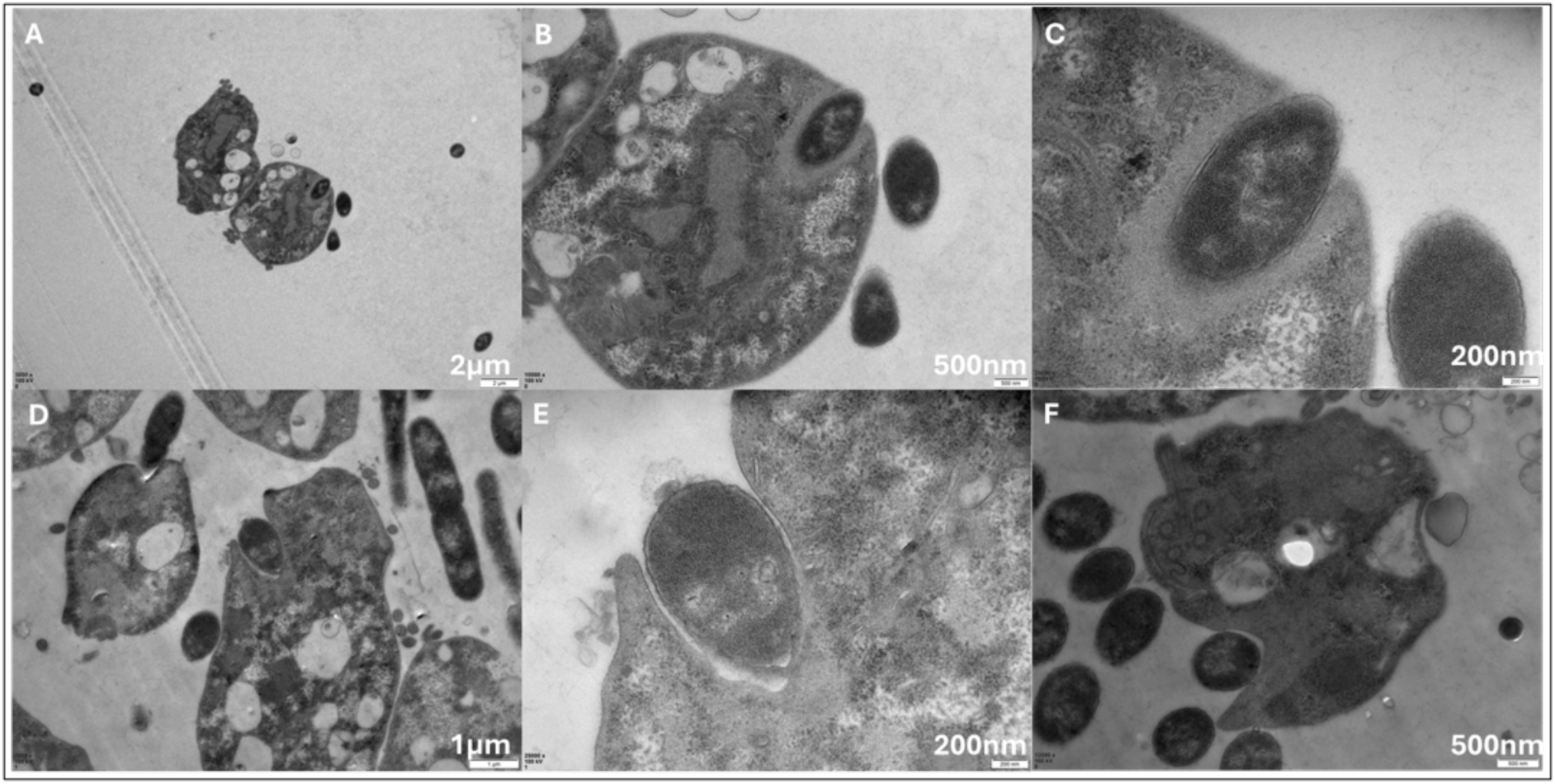
Transmission electron microscopy of *T. gallinae* interacting with *E. coli.* (A) Two *T. gallinae* cells in close proximity to multiple *E. coli* cells (darker cells), including one *E. coli* cell enclosed within a membrane pocket invaginating into a *T. gallinae* cell. (B–C) Higher magnification views of the *T. gallinae* cell and invaginating *E. coli* from panel A, showing progressive detail of the interaction. (D) Multiple *T. gallinae* cells near several *E. coli* cells, with two *E. coli* cells located within distinct invaginating membrane pockets. (E) Higher magnification view of a *T. gallinae* cell from panel D, showing an invaginating *E. coli* cell. (F) A *T. gallinae* cell closely associated with several *E. coli* cells, including one within an invaginating pocket. Scale bars are shown in the bottom right of each panel, with lengths indicated above in white text.

### *Trichomonas gallinae* vastly alters its transcriptome during *in vitro* co-culture with bacteria

The enhanced fitness and motility of *T. gallinae* during bacterial co-culture and the pocket physical interactions suggests *T. gallinae* may modify its transcriptome in response to the presence of *E. coli*. To assess this, total RNA was extracted for RNAseq from the *T. gallinae*-*E. coli* co-cultures and *T. gallinae* monocultures represented in Figure 1A, after 24 hours incubation time in triplicate. The rRNA-depleted raw reads obtained across all six samples, totalling 1,111,396,102 paired reads, are available under Bioproject: PRJNA1019275 at the NCBI’s SRA database (**Table S1**). Reads were selected for good quality (PHRED quality score (Q)> 34) with a base call accuracy of >99.9% for each base pair within a given read (**Figure S4**).

All reads were aligned to the available *T. gallinae* (strain A1, isolate XT-1081/07, clone GF1c) genome (Alrefaei *et al*., 2019) and analysed for differential expression. Principal component analysis revealed a distinct difference in the transcriptomic profiles of monocultured and co-cultured *T. gallinae* (**Figure 3A**). A tagwise biological coefficient of variation plot showed an average biological coefficient of variation (BCV) between 0.2 and 0.5, with a common dispersion of ≈0.29 (**Figure 3B**). The analysis showed that ≈56% of the *T. gallinae* annotated protein-coding genes were significantly (p-value < 0.05) modulated when in co-culture (**Figure 3C**). Of the 21,927 annotated protein-coding genes in the *T. gallinae* genome, 5829 were significantly (p-value < 0.05) upregulated and 6421 were significantly (p-value < 0.05) downregulated in the co-culture condition.

**Figure 3.**
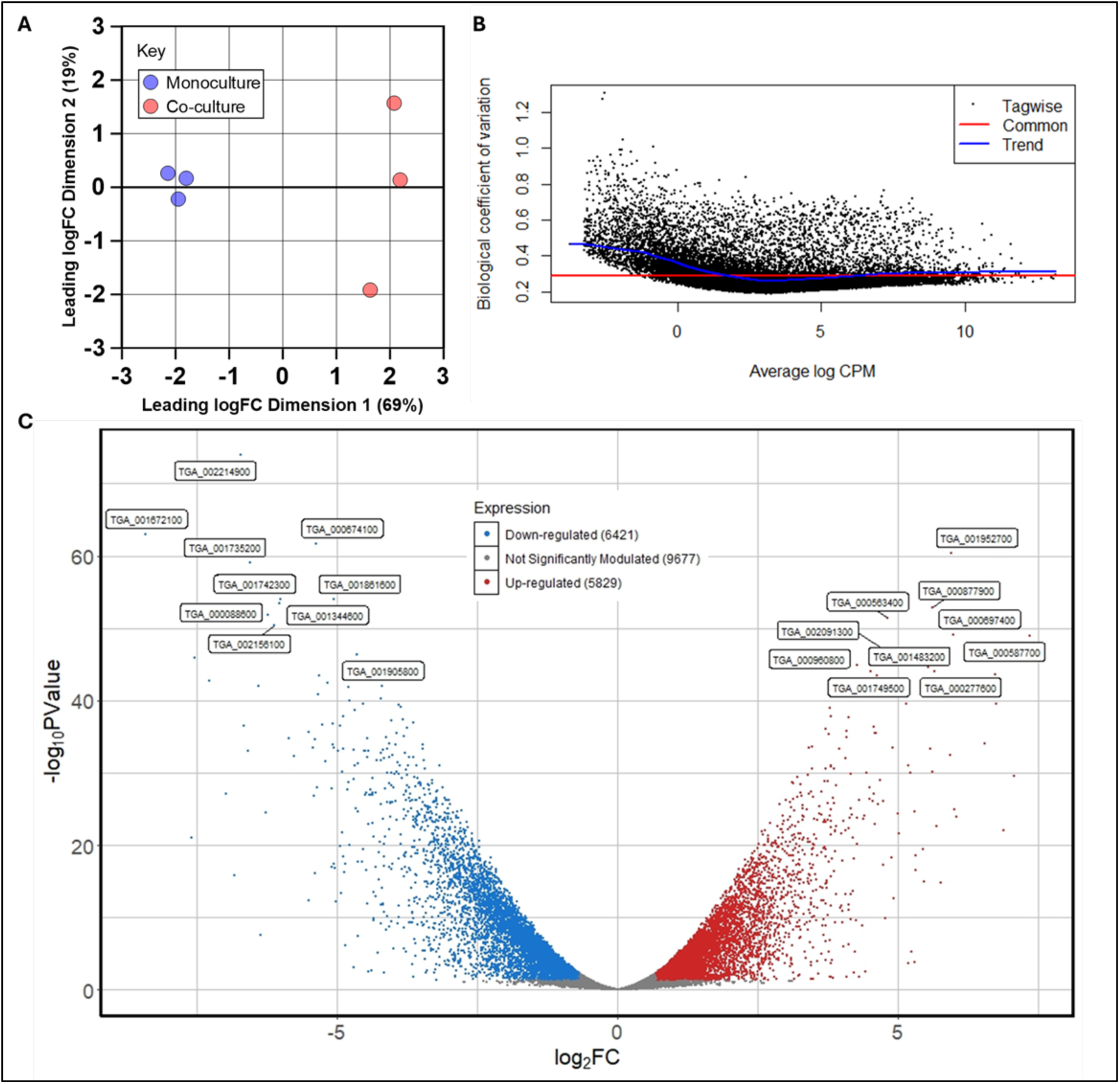
Transcriptomic response of *T. gallinae* to co-culture with *E. coli.* (A) Principal component analysis of *T. gallinae* RNASeq reads from monoculture (blue) and co-culture with *E. coli* (red). (B) Biological coefficient of variation plot showing dispersion estimates for each *T. gallinae* gene relative to average log counts per million (CPM) across monoculture and co-culture conditions. (C) Volcano plot highlighting differential gene expression (log₂ fold change) in *T. gallinae* during co-cultures compared to monocultures. Significantly upregulated genes (p-value < 0.05) are shown in red; significantly downregulated genes (p-value < 0.05) are in blue and those with no significant change (p > 0.05) are in grey. The top 10 most differentially expressed genes are labelled with their gene IDs. Gene annotations along with the full RNAseq data read counts and statistics are provided in Table S4.

Gene ontology (GO) enrichment analysis was performed to identify the functional categories of differentially expressed transcripts. Biological processes such as protein phosphorylation, cellular components like phosphatidylinositol 3-kinase complexes, and multiple molecular functions, including protein serine/threonine kinase activity, were enriched among significantly upregulated *T. gallinae* genes in co-culture (**Figure 4A**). These GO terms are all involved with intracellular signalling consistent with the observed modulation of gene expression during co-culture.

**Figure 4.**
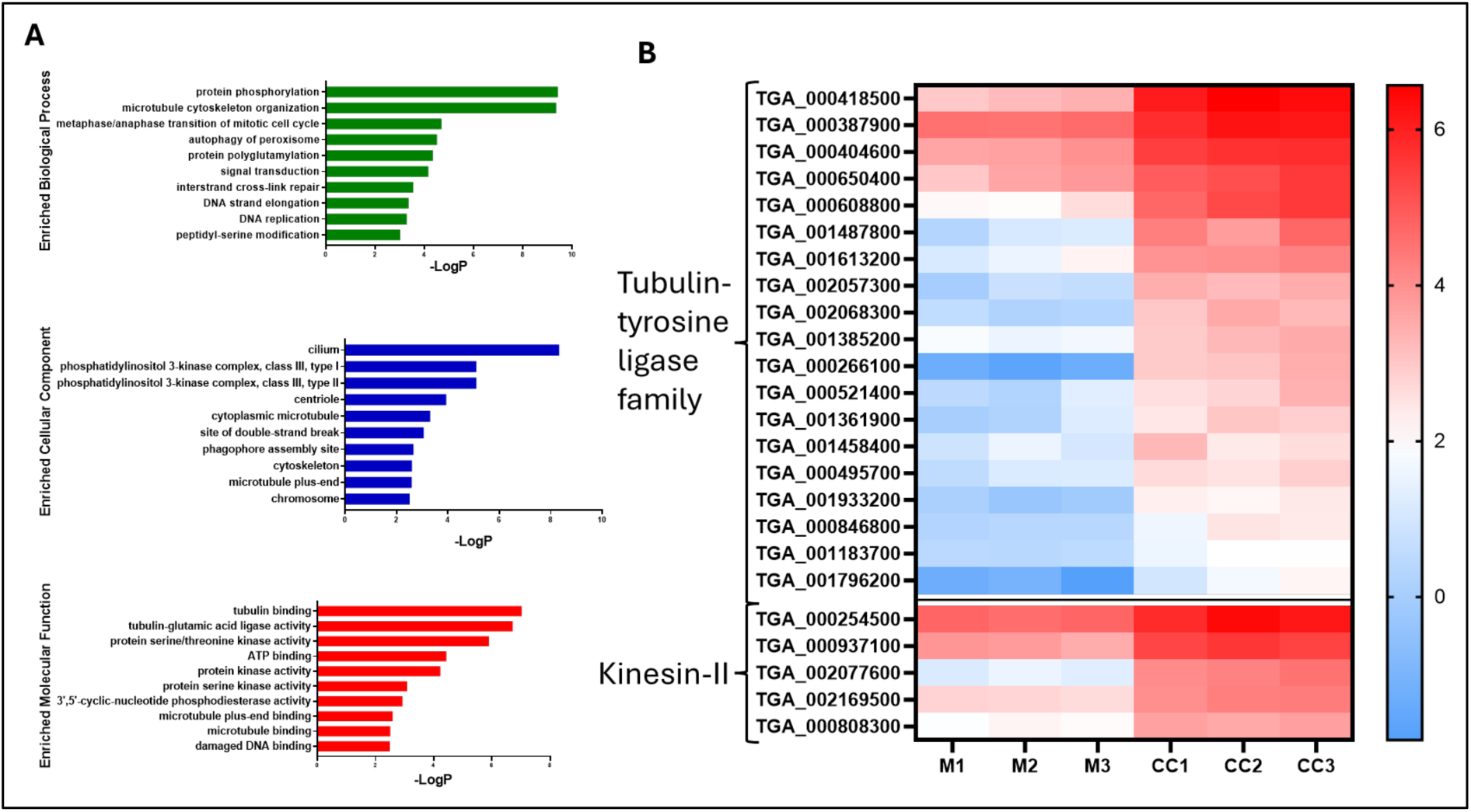
GO enrichment and expression of motility-associated genes in *T. gallinae* during co-culture with *E. coli.* (A) Bar chart showing the top 10 most significantly enriched Gene Ontology (GO) terms in the *T. gallinae*–*E. coli* RNASeq dataset, categorised into biological processes (green), cellular components (blue), and molecular functions (red). Enrichment is represented as –log₁₀(p-value). (B) Heatmap of Log₂TPM values for putative motility-associated genes in *T. gallinae* during co-culture (CC1-3) and monoculture (M1-3), with the associated gene class annotated to the left of each gene’s ID. The colour scale is shown to the right of the heatmap.

To investigate the potential links between the enhanced motility observation based on cell imaging and the significantly modulated transcriptome, we examined GO terms and gene sets involved with motility. Notably, the “tubulin binding” GO category, which includes *T. gallinae* tubulin-tyrosine ligase family proteins associated with flagellar motility (Kubo & Oda, 2017) was enriched among significantly up-regulated transcripts. Additionally, five putative Kinesin-II proteins, also implicated in flagellar motility (Pan & Snell, 2002), were significantly upregulated during co-culture (**Figure 4B**). These significant changes in the transcriptome are consistent with the observed increase in parasite motility (**Figure 1B, C**) in the presence of bacteria.

### *Trichomonas gallinae* expresses candidate bacterial targeting genes when in co-culture with *E. coli*

Given the evidence of modulated *T. gallinae* fitness and physical interactions with *E. coli* during co-culture, RNAseq data was analysed for the expression of genes encoding potential bacterial-targeting enzymes. Our analysis identified modulated genes encoding putative bacterial targeting glycoside hydrolases (GH) and homologues to *T. vaginalis* NlpC/P60 (Barnett *et al*., 2023), suggesting a role in *T. gallinae*–bacteria interactions. One of each structural class of NlpC/P60 endopeptidase, three of five GH19s (putative chitinase/lysozyme), three of four GH25s (putative lysozyme) and one of four GH3s (putative N-acetylglucosaminidase, NagZ-like) had their transcripts significantly upregulated during co-culture. Notably, among the most highly expressed up-regulated genes were one GH19 (TGA_001274700) and a putative AMP (TGA_0001351900) (**Figure 5A**). Therefore, *T. gallinae* significantly upregulated the expression of genes coding for known and putative bacterial PG targeting enzymes and one AMP in the tested co-culture condition.

**Figure 5.**
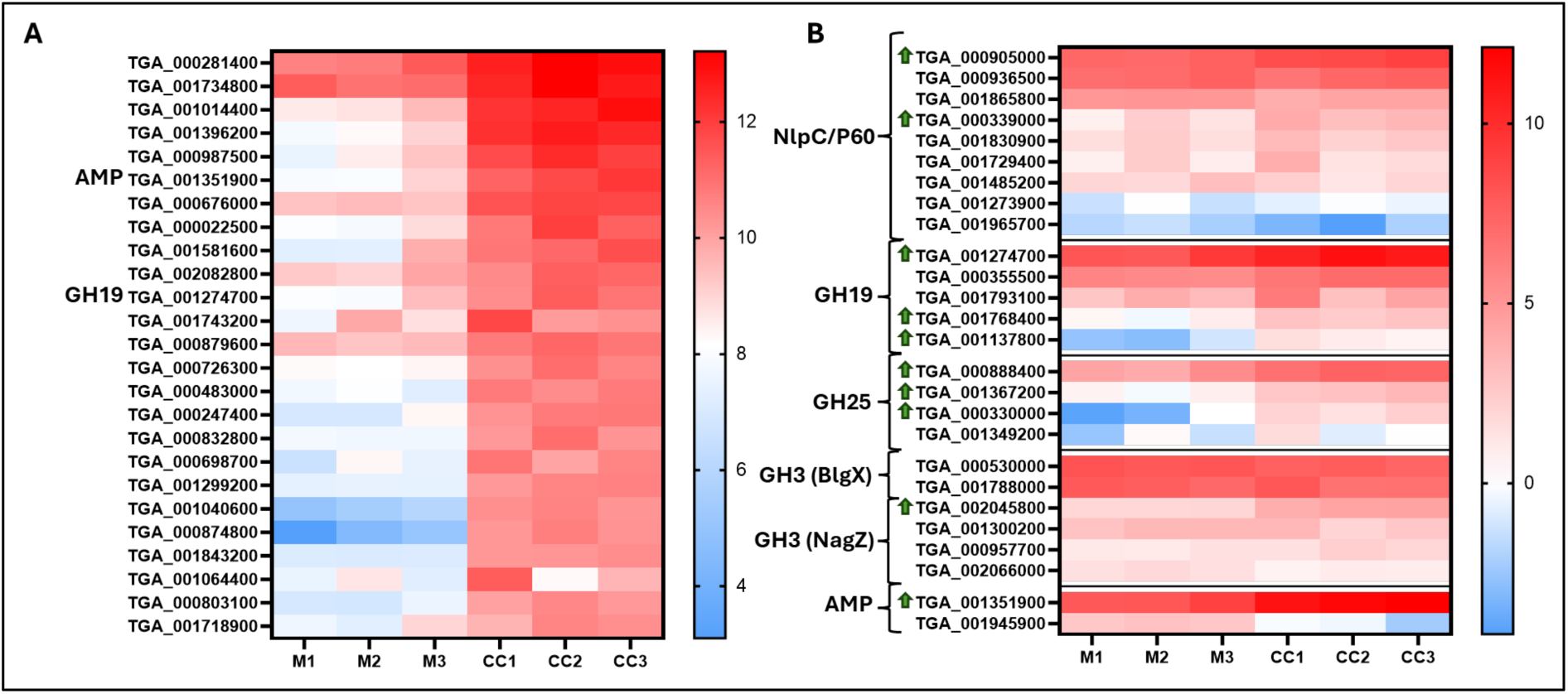
Differential gene expression in *T. gallinae* during co-culture with *E. coli.* (A) Heatmap showing Log₂TPM values of the top 25 most highly expressed and significantly upregulated (p-value < 0.05) genes in *T. gallinae* during co-culture with *E. coli* (CC1-3), alongside their expression levels in *T. gallinae* monoculture (M1-3). Genes encoding members of the antibacterial toolkit are labelled next to their corresponding gene IDs (one GH19 and one AMP-Lcn972). (B) Heatmap of Log₂TPM values for the *T. gallinae* genes encoding antibacterial toolkit components in co-culture (CC1-3) and monoculture (M1-3). Green arrows next to gene IDs indicate significantly upregulated genes (p-value < 0.05), with the associated toolkit class annotated to the left of each gene ID. Colour scales for each heatmap are shown to the right.

### *Trichomonas* GH19s are a conserved group of enzymes laterally acquired from bacteria with distinct structural properties compared with characterised GH19s

To begin the characterization of the putative bacterial-targeting enzymes identified in the co-culture RNAseq, we combined comparative genomics and phylogenetics on *T. gallinae* GH19s. GH19s can be lysozymes or chitinases or, in some circumstances, both (Hoell *et al*., 2006; Wohlkönig *et al*., 2010; Lim *et al*., 2012). Homologues of *T. gallinae* GH19s are present across all sequenced *Trichomonas* species genomes, all belonging to one of two structural domain sub-classes (A and B) (**Figure 6**). Class B GH19s contain two tandem putative PG-binding SH3b domains, absent in Class A. Both classes share conserved putative Glu catalytic residues, predicted through sequence comparison with functionally characterised GH19 enzymes (Hoell *et al*., 2006; Wohlkönig *et al*., 2010; Lim *et al*., 2012). A signal peptide (SP) is predicted to be present in most of the *T. gallinae* GH19s. Due to N-terminal similarity, the presence of a SP is predicted by sequence comparison for all *Trichomonas* GH19s (**Figure S5**). SH3b domains are also present in some *T. vaginalis* NlpC/P60 enzymes (Pinheiro *et al*., 2018; Barnett *et al*., 2023) but are clearly distinct from those identified in the GH19 proteins (**Figure S5**).

**Figure 6.**
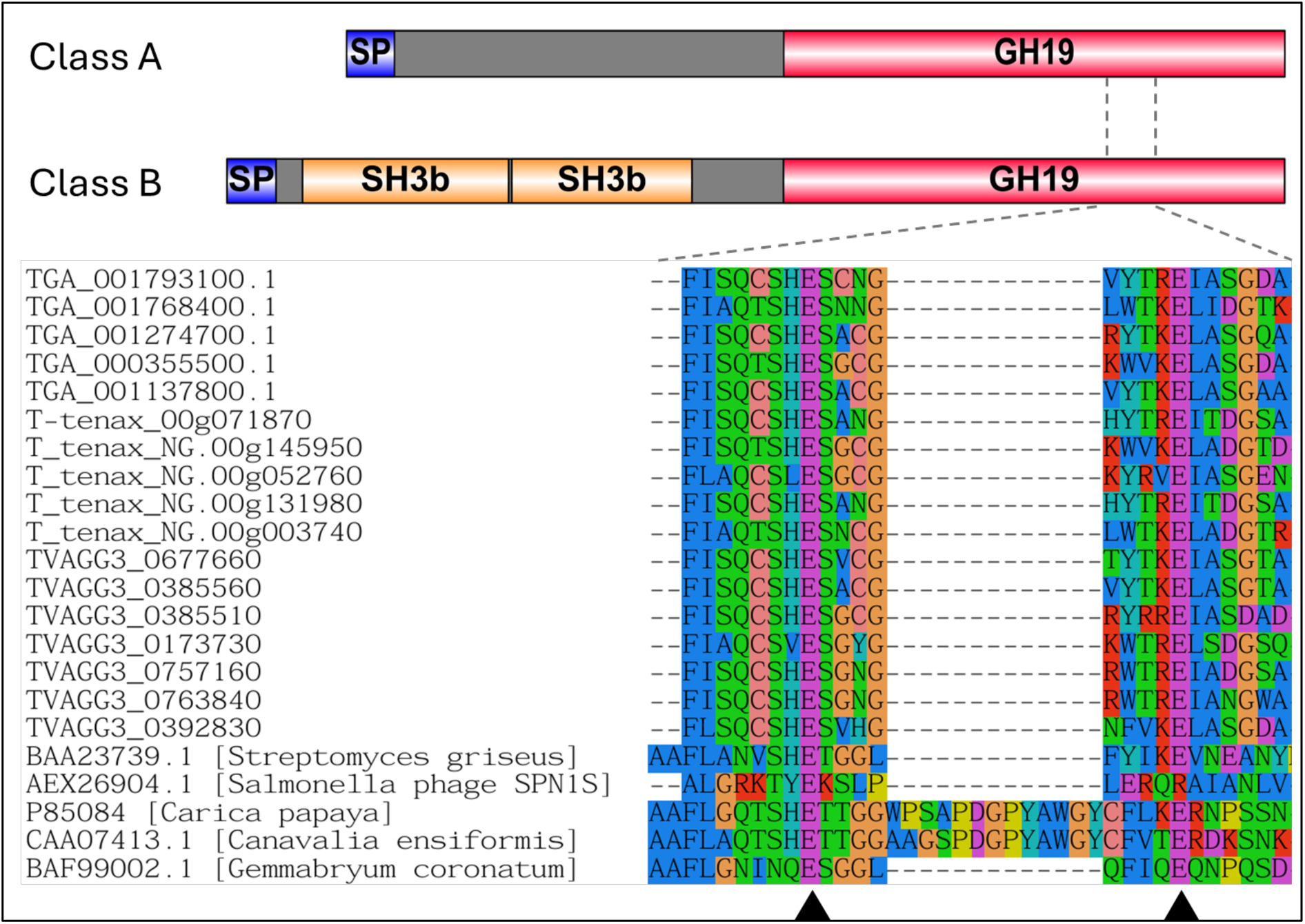
Domain architecture and catalytic site conservation of *Trichomonas* GH19 proteins. Top: Domain organization of Class A and Class B *Trichomonas* GH19 proteins. Signal peptides (SP) are shown in blue, SH3b domains in yellow, and GH19 catalytic domains in red. Grey regions indicate sequence segments without predicted domains. Bottom: Partial multiple sequence alignment of the catalytic region from *Trichomonas* GH19 proteins, aligned with corresponding regions from characterized/reference GH19 enzymes from bacteria, plants, and phage (CAZy database, see main text). Black triangles mark putative catalytic residues in *Trichomonas* sequences and known catalytic sites in reference GH19s. Dashed lines connect the aligned catalytic region to its approximate location within the domain diagrams above.

Phylogenetic analysis indicates that *Trichomonas* GH19s are conserved across sequenced *Trichomonas spp.* and related Trichomonads, *Histomonas meleagridis* and *Tritrichomonas foetus* (**Figure 7 and Figure S6**). Homologues are also present in distantly related metamonads, *Hexamita inflata*, *Monocercomonoides exilis*, and *Blattamonas nauphoetae*. SH3b domains are unique to *Trichomonas spp.* GH19s, while other eukaryotic homologues often feature alternative putative bacterial PG-binding domains, such as LysM. These eukaryotic sequences cluster with a bacterial clade, distinct from CAZy-characterized GH19 sequences, which function as lysozymes, chitinases, or both (**Figure 7**). In conclusion, *Trichomonas* GH19s are a conserved set of likely functional proteins structurally distinct from characterized GH19s in the CAZy database. Likely acquired via LGT from bacteria, they have since diversified within Metamonada, incorporating additional putative PG-binding domains through potential separate LGT events and domain reshufling.

**Figure 7.**
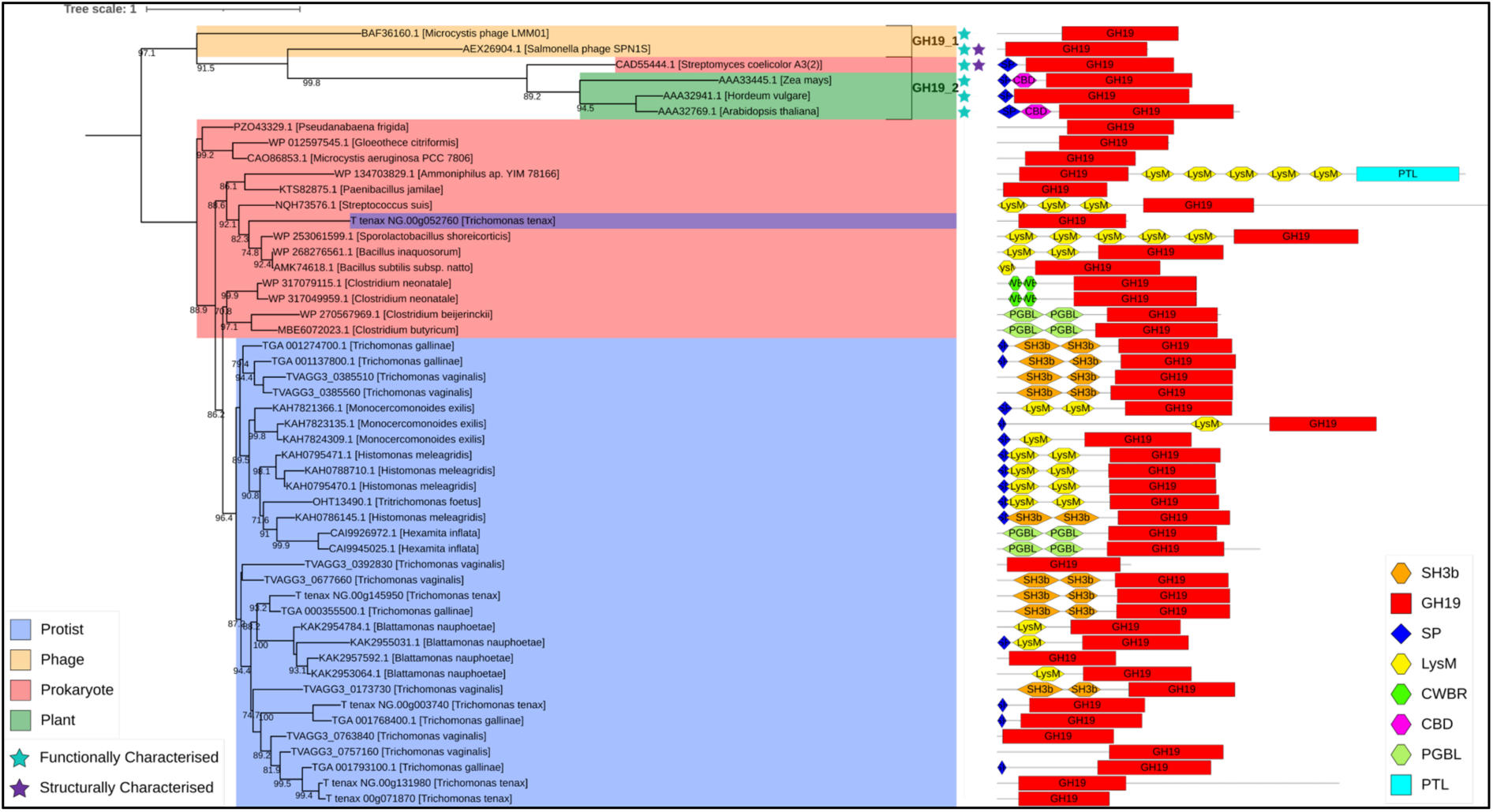
Phylogenetic analysis and domain architecture of GH19 proteins. Outgroup-rooted phylogenetic tree constructed from a masked multiple sequence alignment of *Trichomonas* and Trichomonad GH19 proteins, homologues identified via BLASTp searches against the NCBI’s nr database, and characterised GH19s from the CAZy database. The phylogeny was generated using the LG+G4 model with ultrafast bootstrapping; bootstrap values >70 are shown at the corresponding nodes. Branch labels include gene ID and species of origin. Clades are colour-coded by taxonomic group: protists (blue), phages (yellow), prokaryotes (red), and plants (green). CAZy classifications of characterised GH19s are noted to the right of relevant leaves. Functionally characterised proteins are marked with a light blue star (top of the figure), and structurally characterised proteins with a purple star. Domain architectures are displayed to the right of each branch, with domains annotated according to the key in the bottom right of the figure.

### *Trichomonas* GH19s are muramidases and not chitinases

*Trichomonas spp.* GH19s are predicted to have either lysozyme (muramidase) or chitinase activity. To test these predictions, we characterised recombinant *T. gallinae* (TGA_001274700.1) and *T. vaginalis* (TVAGG3_0677660) biochemically. For TGA_001274700.1 we also generated an inactivity mutant (E202Q) (Taira *et al*., 2011).

At first, proteins were tested using a commercial lysozyme activity kit (Sigma Aldrich, LY0100) (**Figure S7**). This kit uses *Micrococcus luteus* cell lysis as a proxy for lysozyme-like activity (Shugar, 1952). Both tested GH19 proteins displayed lysozyme-like activity compared to their respective controls.

PG is the canonical substrate for lysozyme as it cleaves the β1,4-glycosidic bond between the N-acetylmuramic acid (Mur*N*Ac) and N-acetylglucosamine (Glc*N*Ac) residues in PG (Vollmer *et al*., 2008). Next, we tested if TGA_001274700.1 and TVAGG3_0677660 can degrade PG purified from *E. coli* BW25113 or MC1061, analysing digestion products by high-performance liquid chromatography (HPLC) (Glauner, 1988) (**Figure 8**). As a positive control we used the muramidase cellosyl, a GH25 from *Streptomyces coelicolor* (Rau *et al*., 2001). Both WT *Trichomonas* GH19s were able to produce similar degradation patterns compared to the control (**Figure 8**). We then analysed the most prominent HPLC peaks produced from digestion of PG purified from *E. coli* MC1061 to confirm, by mass spectrometry, that degradation products indeed originated from PG and matched known muramidase reaction products (**Figure S8**). Considering the bioinformatic prediction in the first place and the presented biochemical evidence, we can conclude that, both TGA_001274700.1 and TVAGG3_0677660 are muramidases with lysozyme-like activity.

**Figure 8.**
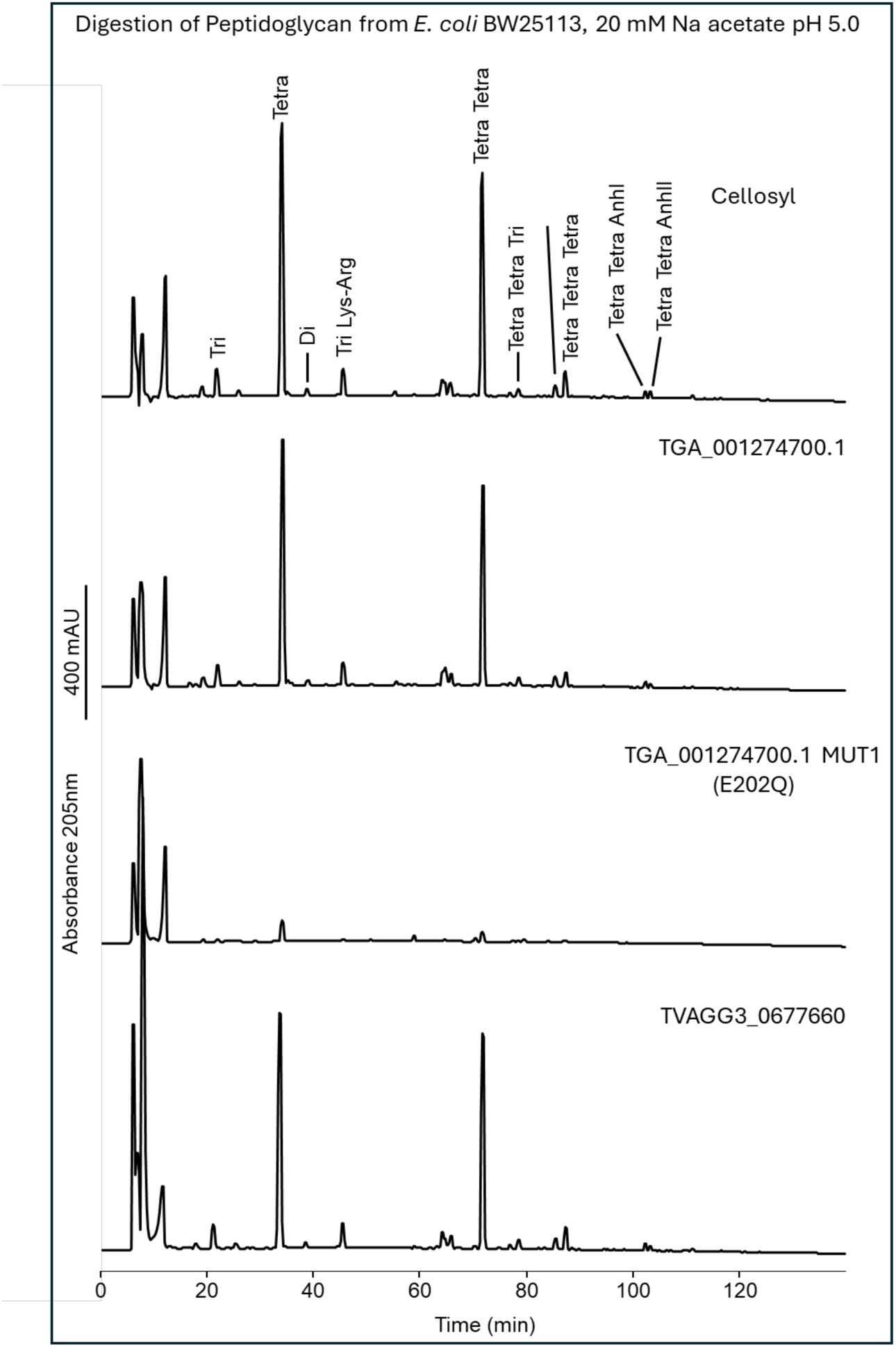
HPLC analysis of peptidoglycan degradation by *Trichomonas* GH19 proteins. Stacked HPLC chromatograms showing peptidoglycan degradation products generated in 20 mM sodium acetate buffer (pH 5.0). From top to bottom: Cellosyl (positive control, a GH25 from *Streptomyces coelicolor*), WT recombinant *T. gallinae* GH19 (TGA_001274700.1), mutant *T. gallinae* GH19 (TGA_001274700.1 MUT1 [E202Q]), and WT recombinant *T. vaginalis* GH19 (TVAGG3_0677660). Major peaks corresponding to muropeptides are labelled only in the cellosyl chromatogram (top panel). All profiles except for the E202Q mutant *T. gallinae* GH19 show a typical lysozyme digestion pattern. The scale bar (400 mAU) is shown to the left of all chromatograms.

GH19 family proteins may also be chitinases, as some can cleave the β1-4 glycosidic bond of various polymers of Glc*N*Ac that are present in chitin (Hamid *et al*., 2013). We therefore tested both wild type (WT) GH19 proteins for their activity against chitin using a commercial chitinase enzymatic assay. The assay contains three structurally different chitin substrates, which allow for testing various types of chitinolytic activity *in vitro*. TGA_001274700.1 and TVAGG3_0677660 were unable to degrade any of the chitin variants compared to the positive control (**Figure S9**). Our result suggests that both enzymes are not chitinases.

### *Trichomonas* has laterally acquired NagZ-like GH3s that could also be active on peptidoglycan

GH3s have a large range of characterised functions (33), including N-acetylglucosaminidase (Cantarel *et al*., 2009). One of six *T. gallinae* GH3s was significantly upregulated during co-culture (**Figure 5**). Therefore, to further characterise the putative bacterial-targeting enzymes identified by comparative RNAseq, we conducted comparative genomics and phylogenetics on *T. gallinae* GH3s to assess the potential function and relevance for *Trichomonas-*bacteria interactions.

GH3 sequences are conserved across sequenced *Trichomonas spp.*, and like GH19, homologues are found in distantly related Trichomonads, *Histomonas meleagridis* and *Tritrichomonas foetus* (**Figure 9 and Figure S10**). The phylogenetic analyses recovered the *Trichomonas* GH3s cluster within a large bacterial clade made up of putative or characterised NagZ (PG targeting β-N-acetylglucosaminidase) enzymes. An outgroup of the *Trichomonas* GH3s and the Trichomonad homologues cluster separately with alternative bacterial homologues. The outgroup *Trichomonas* GH3s are similar (determined with the use of Foldseek using Alphafold predictions of the outgroup *Trichomonas* GH3s (Jumper *et al*., 2021; van Kempen *et al*., 2024)) to another bacterial GH3 – BglX (a β-glucosidase, also a PG targeting enzyme). Thus, *Trichomonas* likely acquired bacterial GH3 enzymes through two separate events: one specific to *Trichomonas spp*., encoding NagZ-like proteins, and another, conserved in Metamonads and *Trichomonas spp.*, encoding BglX-like proteins.

**Figure 9.**
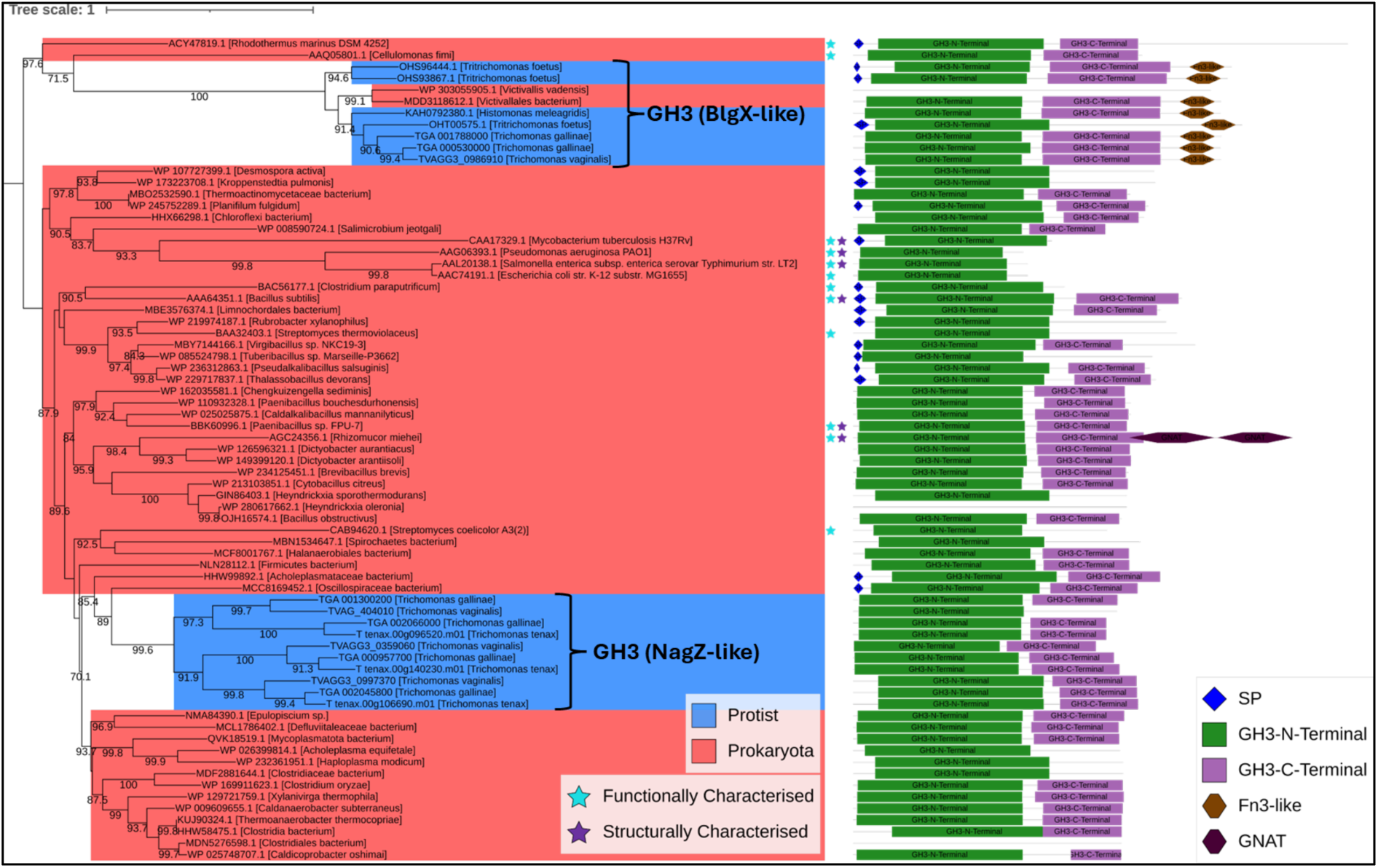
Phylogenetic analysis and domain architecture of GH3 proteins. Outgroup-rooted phylogenetic tree generated from a masked multiple sequence alignment of *Trichomonas* and trichomonad GH3 proteins, homologues identified via BLASTp searches against the NCBI’s nr database and characterised GH3S from the CAZy database. The maximum likelihood phylogeny was inferred using the LG+I+G4 model with ultrafast bootstrapping; bootstrap values >70 are shown at the corresponding nodes. The scale bar shows the inferred number of substitutions per site. Branch labels include gene ID and species of origin. *Trichomonas* NagZ-like and BlgX-like GH3s are indicated to the right of relevant branches. Clades are colour-coded by taxonomic group: protists (blue) and prokaryotes (red). Functionally characterised proteins are marked with light blue stars; structurally characterised proteins with purple stars. Domain architectures are shown to the right of each branch, with domains annotated as per the key in the bottom right of the figure. Three proteins did not have hits for the indicated domain profiles despite being in the Blast hit list.

NagZ, in *E. coli*, is responsible for the cleavage of an intermediate within the PG recycling pathway Glc*N*Ac-1,6-anhydro*M*urNAc(peptide) (Vötsch & Templin, 2000). To assess the functionality of *Trichomonas* NagZ-like GH3s, sequence analysis was performed. All possess the NagZ consensus motif and the two key catalytic residues (histidine and aspartic acid) (e.g. TGA_000530000.1 His186 Asp 264) (Bacik *et al*., 2012) (**Figure 10**). They share a conserved domain organization, featuring N and C-terminal GH3 domains, a cytoplasmic domain, and a transmembrane domain. These findings suggest *Trichomonas* NagZ-like GH3s function as glucosaminidases, acting on whole or partially digested PG. However, this could not be tested experimentally as attempts to produce soluble recombinant GH3 proteins from *T. gallinae* (TGA_002045800.1) and *T. vaginalis* (TVAGG3_0997370) all failed.

**Figure 10:**
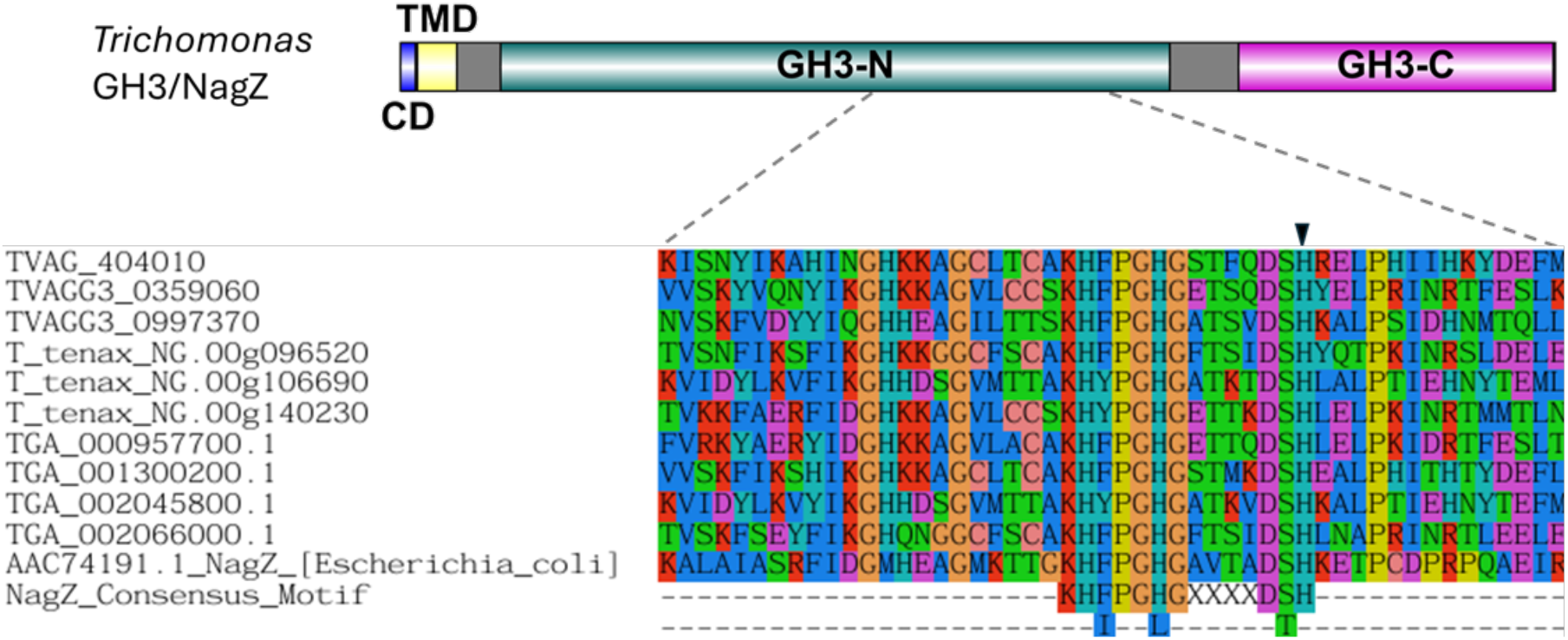
Domain architecture and consensus motif conservation of *Trichomonas* GH3 (NagZ-like) proteins. Top: Domain organization of *Trichomonas* GH3 (NagZ-like) proteins. Cytoplasmic domains (CD) are shown in blue, transmembrane domains (TMD) in yellow, and GH3 N-terminal (GH3-N) and C-terminal (GH3-C) catalytic domains are in green and purple, respectively. Bottom: Partial multiple sequence alignment of the NagZ consensus motif region from *Trichomonas* GH3 proteins aligned with the corresponding region of *E. coli* NagZ. The black triangle indicates one of the two putative catalytic residues in *Trichomonas* sequences and the known corresponding catalytic residue in *E. coli* NagZ (see main text). The NagZ consensus motif is shown at the bottom of the alignment (see main text) over two lines to indicate variations in the motif. Dashed lines connect the aligned catalytic region to its corresponding location in the domain diagram above.

### *Trichomonas* can use peptidoglycan breakdown products as metabolites

*Trichomonas spp.* have functional muramidases (**Figure 8**) and likely functional NagZ-like enzymes (**Figure 10**). Therefore, *Trichomonas spp.* are likely able to hydrolyse PG and release free Glc*N*Ac, which is a potential source of carbon and nitrate. Using Glc*N*Ac as a metabolite could contribute to the observed enhanced fitness (**Figure 1A**) and motility (**Figure 1B**) of *T. gallinae* in co-culture with *E. coli*. To test if Glc*N*Ac could be utilised we identified the putative catabolic pathway to commit Glc*N*Ac to glycolysis in *T. gallinae* (**Figure 11C**).

**Figure 11:**
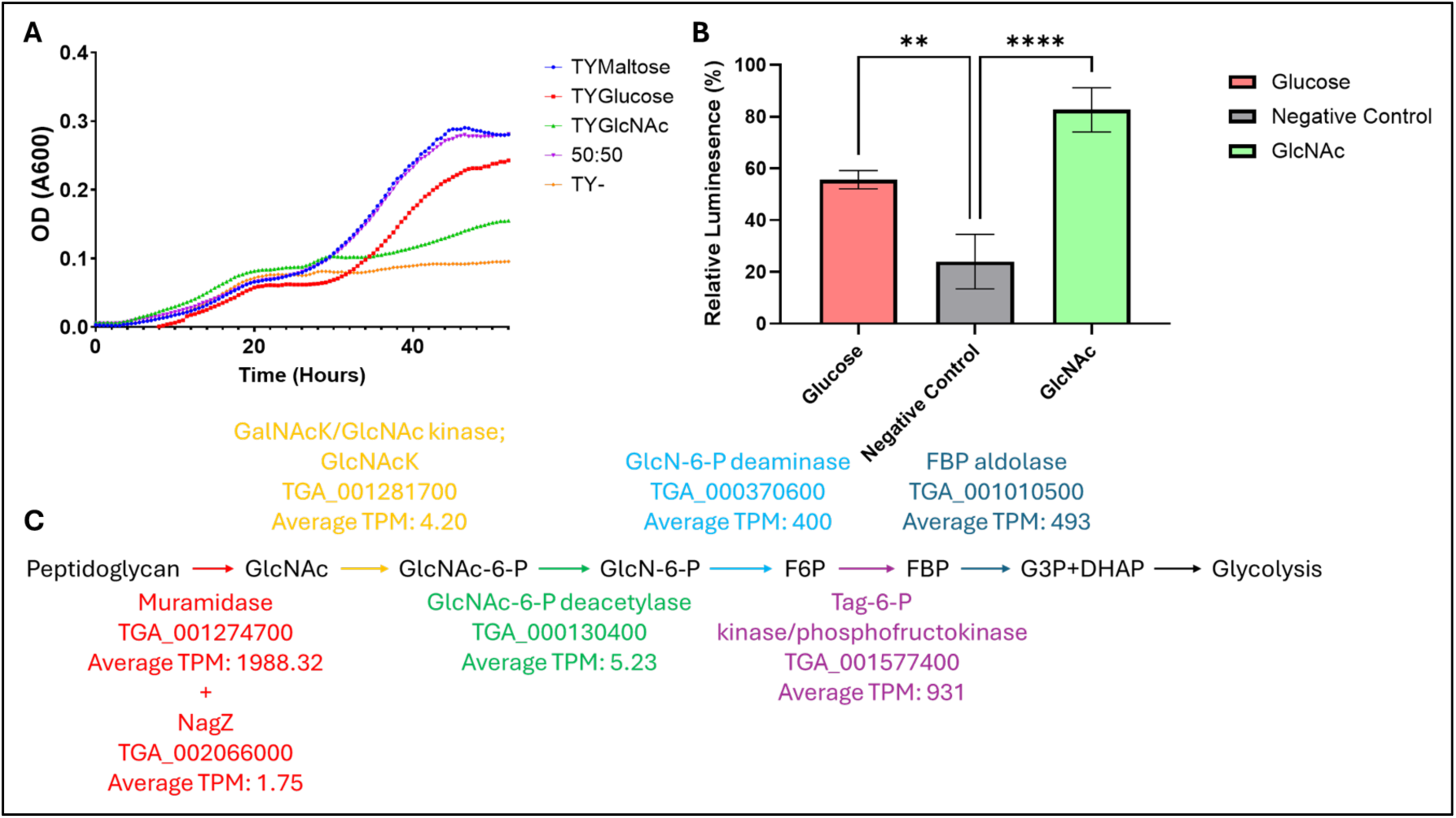
Utilization of Glc*N*Ac by *T. gallinae.* (A) Growth curve showing the mean OD (A600) of *T. gallinae* cultured in standard medium (TYMaltose, blue), maltose-free medium (TY–, yellow), and media supplemented with glucose (TYGlucose, red), Glc*N*Ac (TYGlc*N*Ac, green), or a 50:50 mixture of glucose and Glc*N*Ac (purple), over 48 hours at 37°C (n = 8). (B) Bar chart showing relative luminescence (%) from an ATP-based viability assay of starved *T. gallinae* cells reintroduced to a medium containing glucose (positive control, red), Glc*N*Ac (green), or without any added monosaccharide,(negative control, grey) medium (n = 3). Asterisks denote significant differences based on one-way ANOVA: ** p-value < 0.005, **** p-value < 0.0001. (C) Schematic of the predicted metabolic pathway by which *T. gallinae* channels Glc*N*Ac (derived from peptidoglycan) into glycolysis. Metabolites are labelled in black; coloured arrows trace the enzymatic steps of the pathway. Arrow colours correspond to the *T. gallinae* gene predicted to encode the enzyme for each step, with average TPM values for these genes (during *T. gallinae*–*E. coli* co-culture) shown in matching colours.

To assess whether *T. gallinae* could utilise Glc*N*Ac as a carbon source *in vitro*, *T. gallinae* was grown in standard TYM medium or standard *Trichomonas spp.* medium with the maltose removed and then supplemented with a range of alternative carbon sources, including Glc*N*Ac. *T. gallinae* grew in TY + Glc*N*Ac but reached a higher maximal OD in TYM (**Figure 11A**).

To determine whether Glc*N*Ac was committed to ATP production in the absence of other metabolites, a luminescence-based ATP assay was performed using various monosaccharides. Ribose, Glucose, GalNAc, and Glc*N*Ac supported significantly higher ATP production levels in *T. gallinae* compared to the negative control after starvation (**Figure 11B**). This further supports the idea that *T. gallinae* can utilize Glc*N*Ac for ATP generation and ultimately growth. Following starvation conditions, it produces more ATP using Glc*N*Ac alone than with more common cell culture monosaccharides like glucose, aligning with the presence of the predicted glycolysis pathway. Overall, PG breakdown products, generated by the combined action of a muramidase and NagZ-like enzyme/s, can serve as an ATP source for *T. gallinae*, which could contribute to the observed fitness advantage in bacterial co-cultures.

### *Trichomonas* have conserved laterally acquired anti-microbial peptides

Among the most transcribed, significantly upregulated genes observed during *T. gallinae*–*E. coli* co-culture RNAseq was TGA_001351900 (**Figure 5A**). This gene encodes a protein containing a Lac_972 domain (InterPro accession IPR006540), also found in the bacterial AMP lactococcin 972 (AMP-Lcn972), which targets the bacterial cell wall precursor Lipid II (Martínez *et al*., 1996; Martínez *et al*., 2008).

Homology searches identified a paralogue of TGA_001351900 in the *T. gallinae* genome (TGA_001945900) and two homologues in the sequenced genome of *T. vaginalis* and *T. tenax*. All six possess a C-terminal Lac972 domain (**Figure 12**), with TGA_001351900 and T_tenax_NG.00g044980 also featuring a predicted signal peptide (**Figure S11**). TGA_001945900 appears incomplete due to a missing start codon. Using each *Trichomonas* AMP-Lcn972 as a BLASTp query against a structurally characterized lactococcin 972 AMP (accession MCT0507501.1), revealed that only three *Trichomonas* AMPs share significant similarity with lactococcin 972 from *Lactococcus cremoris* (**Table 1**). However, all six are predicted to be antimicrobial peptides according to the CAMPR3 AMP prediction with an SVM classifier (Waghu & Idicula-Thomas, 2020). Broader homology searching using *Trichomonas* AMP-Lcn972 as a query found homologous sequences from other metamonads: *Tritrichomonas foetus*, *Histomonas meleagridis*, *Hexamita inflata*, *Monocercomonoides exilis* and *Streblomastix strix* and several gram-positive bacterial genera (**Table S2**). Phylogenetic analysis indicates that *Trichomonas* and metamonad AMP sequences resemble bacterial sequences but cluster separately from the characterized lactococcin 972 of *Lactococcus cremoris* (**Figure 13**). Sequence alignment reveals a putative motif near two fully conserved histidine residues, not previously described in lactococcin 972 AMPs. AlphaFold2 predictions reveal high structural similarity between *Trichomonas* AMPs and MCT0507501.1 from *Lactococcus cremoris* (**Figure 14**). Overall, bioinformatic investigations reveal a conserved set of putative AMPs within *Trichomonas* and Metamonads, which are similar to a known AMP and are therefore likely functional.

**Figure 12:**
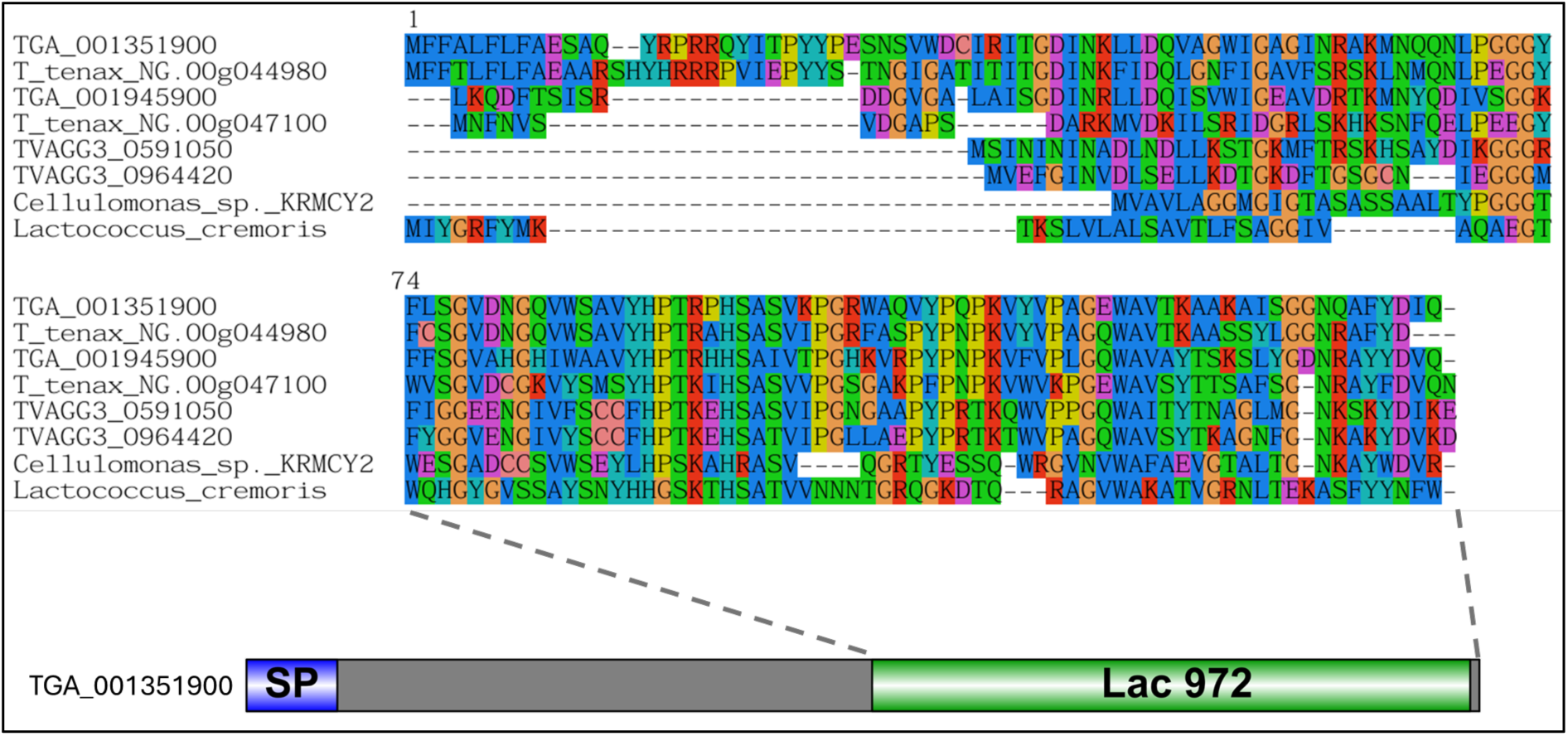
Domain architecture and sequence conservation of *Trichomonas* AMP-Lcn972. Top: Multiple sequence alignment of *Trichomonas* antimicrobial peptides (AMPs) aligned with *Lactococcus cremoris* and *Cellulomonas sp.* KRMCY2 Lcn972. Bottom: Domain organisation of *Trichomonas* AMP-Lcn972, with predicted signal peptide (SP) shown in blue and the Lac972 domain in green. Dashed lines link the aligned region to its corresponding location in the domain diagram below.

**Table 1.** Selected features of AMP-Lcn972 from *Trichomonas gallinae, T. vaginalis* and *T. tenax*.

| Gene_ID | Species | Pseudonym | Length (AA) | GRAVY | Predicted SP | BLASTp bitscore vs MCT0507501.1 | BLASTp Evalve vs MCT0507501.1 | BLASTp %Identity vs MCT0507501.1 | CAMPR3 AMP prediction (SVM classifier) |
| --- | --- | --- | --- | --- | --- | --- | --- | --- | --- |
| TVAGG3_0591050 | *Trichomonas vaginalis* | vAMP1 | 103 | -0.48932 | No | N/A | N/A | N/A | 1.000 |
| TVAGG3_0964420 | *Trichomonas vaginalis* | vAMP2 | 99 | -0.2596 | No | 24.6 | 6.00E-05 | 28.81% | 0.997 |
| TGA_001351900.1 | *Trichomonas gallinae* | gAMP1 | 137 | -0.32482 | Yes | N/A | N/A | N/A | 1.000 |
| TGA_001945900.1 | *Trichomonas gallinae* | N/A | 120 | -0.27143 | No | 18.9 | 0.010 | 26.00% | 1.000 |
| TGA_001945900.2 | *Trichomonas gallinae* | gAMP2 | 77 | -0.35584 | No | 19.6 | 0.003 | 26.00% | 0.043 |
| T_tenax_NG.00g044980.m01 | *Trichomonas tenax* | tAMP1 | 136 | -0.15882 | Yes | N/A | N/A | N/A | 1.000 |
| T_tenax_NG.00g047100.m01 | *Trichomonas tenax* | tAMP2 | 110 | -0.49545 | No | 31.2 | 3.00E-07 | 36.73% | 1.000 |

**Figure 13:**
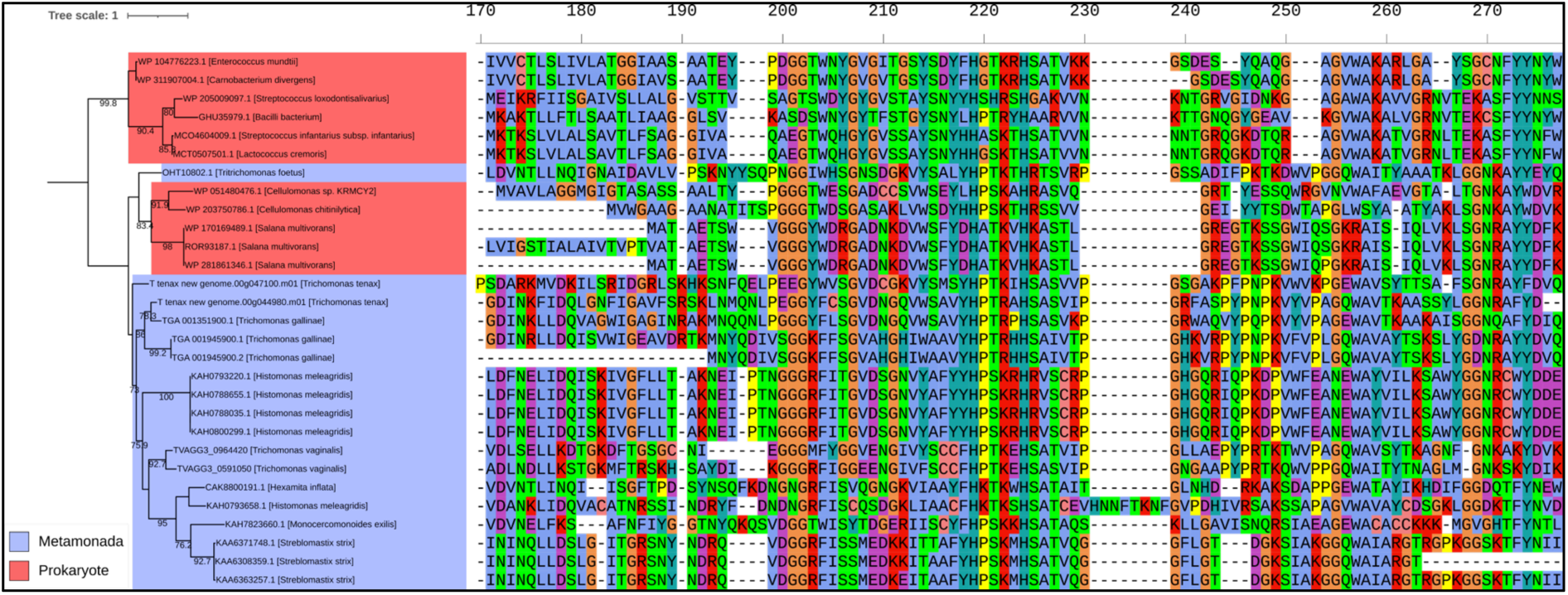
Phylogenetic analysis and multiple sequence alignment of selected AMP-Lcn972 proteins. Outgroup-rooted phylogenetic tree generated from a masked multiple sequence alignment of *Trichomonas* and trichomonad AMP-Lcn972 proteins, along with homologues identified via BLASTp searches against the NCBI’s nr database. The maximum likelihood phylogeny was inferred using the WAG+G4 model with ultrafast bootstrapping; bootstrap values >70 are shown at the corresponding nodes. The scale bar shows the inferred number of substitutions per site. Branches’ labels include gene ID and species of origin. Clades are colour-coded by taxonomic group: Metamonada (blue) and prokaryotes (red). To the right of the tree, a partial section of the multiple sequence alignment highlights conserved residues, illustrating sequence conservation across the proteins used in the phylogenetic analysis.

**Figure 14:**
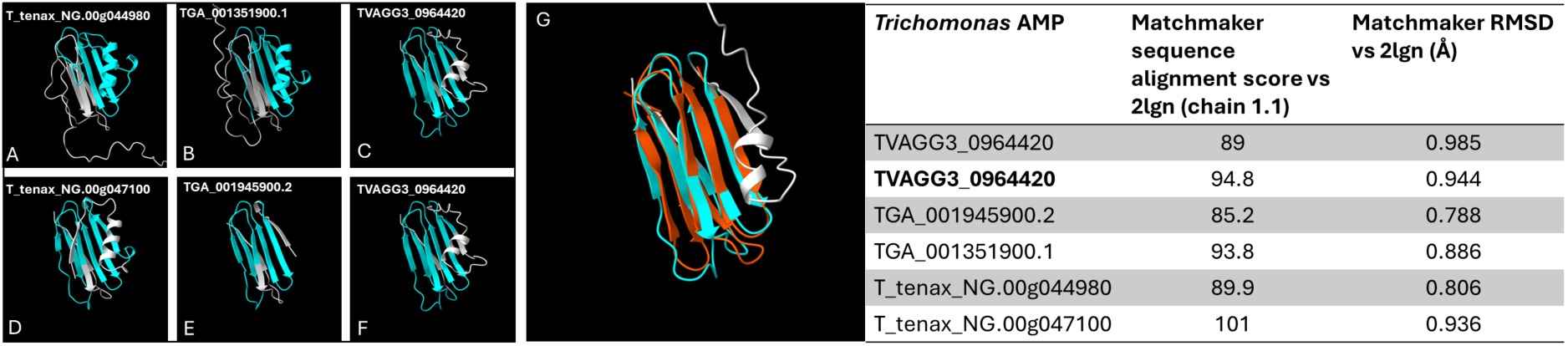
AlphaFold2 predicted structures of *Trichomonas* AMP-Lcn972s and overlay with Lcn972 from *Lactococcus lactis* (PDB 2lgn), with Matchmaker summary table. (A-F) AlphaFold2 predicted structures of each *Trichomonas* AMP-Lcn972, with the corresponding gene ID labelled above each structure. Structures are coloured, with the predicted Lac972 domain in blue. (G) Overlay of TVAGG3_0964420 with 2lgn, where the predicted Lac972 domain of TVAGG3_0964420 is shown in blue and the Lac972 domain of 2lgn in orange. To the right of all structures is a table summarising the Matchmaker output for all *Trichomonas* AMP-Lcn972s vs 2lgn.

To evaluate *Trichomonas* AMP-Lcn972 functionality, *E. coli* transformed with plasmids encoding *Trichomonas* recombinant AMP-Lcn972 (the conserved domain) were generated for expression and purification. Most transformed *E. coli* exhibited growth defects post-induction (**Figure S12**). Minimal soluble recombinant AMP was recovered from cell lysis (**Figure S13**). These data are consistent with these recombinant AMP-Lcn972s being toxic to *E. coli*.

### *Trichomonas* GH25s represent another group of conserved laterally acquired putative muramidase enzymes

During *T. gallinae*–*E. coli* co-culture RNAseq, GH25-coding genes were upregulated, with three of five significantly so. Known GH25 enzymes function exclusively as muramidases (Cantarel *et al*., 2009). Blast searches and phylogenetic analysis suggest GH25 sequences are conserved in *Trichomonas spp.*, likely acquired via LGT from prokaryotes (**Figure 15**). Additionally, *Trichomonas* GH25s retain key catalytic residues for muramidase activity (**Figure 15**). Thus, *Trichomonas* GH25s represent another conserved, laterally acquired group of PG-targeting enzymes.

**Figure 15.**
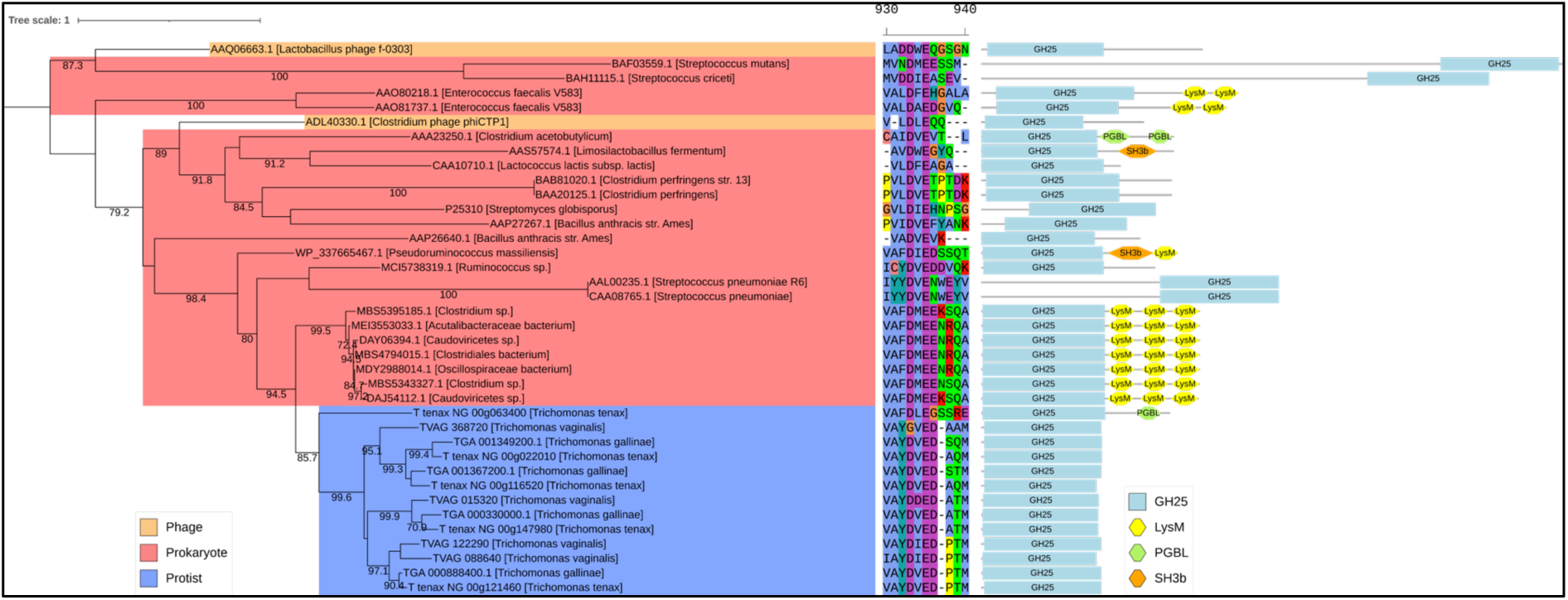
Phylogenetic analysis and domain architecture of GH25 proteins. Midpoint-rooted phylogenetic tree generated from a masked multiple sequence alignment of *Trichomonas* GH25 proteins, homologues identified via BLASTp searches against the NCBI’s nr database, and selected characterised GH25s from the CAZy database. The maximum likelihood phylogeny was inferred using the LG+I+G4 model with ultrafast bootstrapping; bootstrap values >70 are shown at the corresponding nodes. The scale bar shows the inferred number of substitutions per site. Branch labels include gene ID and species of origin. Clades are colour-coded by taxonomic group: protists (blue), phage (orange), and prokaryotes (red). In the centre is a partial segment of the multiple sequence alignment used to generate the phylogeny illustrating the conserved GH25 catalytic region. Domain architectures are displayed to the right of each branch, with domains annotated according to the key in the bottom right of the figure.

## Discussion

Emerging evidence indicates that the outcome of *Trichomonas*-host interactions is significantly influenced by the composition of associated bacterial communities. In *T. vaginalis*, the most extensively characterized species, distinct bacterial partners have been shown to modulate both parasite-mediated cytopathic effects and host inflammatory responses. These synergistic interactions are considered central to the epithelial disruption and mucosal damage that define the pathobiology of trichomoniasis (Mercer & Johnson, 2018; Riestra *et al*., 2022). A striking case involves two cell wall-deficient, mycoplasma-like bacterial species that frequently co-occur with *T. vaginalis* infections as endosymbionts (Fettweis *et al*., 2014; Margarita *et al*., 2022), which have been shown to significantly alter the parasite’s transcriptomic profile, both individually and in combination. These transcriptomic changes are associated with enhanced virulence phenotypes, including increased adherence to human epithelial cells, and heightened haemolytic activity (Margarita *et al*., 2022). In contrast, the molecular mechanisms underlying *Trichomonas* interactions with bacteria possessing a cell wall remain poorly characterized. A notable exception is the NlpC/P60 family of peptidases, which have been experimentally shown in *T. vaginalis* to degrade PG from both Gram-negative (*E. coli*) and Gram-positive (*L. gasseri*) bacteria (Pinheiro *et al*., 2018; Barnett *et al*., 2023). Expression of these NlpC/P60 encoding genes at the mRNA level is significantly upregulated in response to exposure to either bacteria, and homologues for these genes were also encoded by the genomes of *T. gallinae* and *T. tenax,* further supporting important roles for these PG hydrolases in mediating parasite-bacteria interactions (Barnett *et al*., 2023). However, beyond these findings, the molecular basis of *Trichomonas* interactions with wall-containing bacteria, and the extent to which such interactions might influence the pathobiology of distinct *Trichomonas* species across different hosts (mammals and birds), remains largely unexplored. To address some of these limitations, we established a simple co-culture model system between the pigeon infecting *T. gallinae* and *E. coli* to investigate through comparative transcriptomics and genomics the protein complements of *Trichomonas* species deployed to target cell wall-containing bacteria.

Testing different ratios of *T. gallinae*/*E. coli* initial cell densities, established a 10:1 ratio as a condition that ensured that *T. gallinae* cell density was nearing a maximal value at the same time as *E. coli* (**Figure 1**). This accounted for *E. coli*’s higher multiplication rate compared to *T. gallinae* and the assumption that *E. coli* more efficiently absorbs nutrients from TYM media via diffusion due to its higher surface-area-to-volume ratio. Despite a lower maximal density, *T. gallinae* can grow at a generally usual rate during co-culture with *E. coli*, but notably persists longer than a monocultured counterpart. This suggests that the presence of *E. coli* may represent a reservoir of nutrients that *T. gallinae* can exploit. Yet it has been described that *T. vaginalis* cannot phagocytose *E. coli*, unlike some other bacterial species (Francioli *et al*., 1983; Juliano *et al*., 1991). This observation was consistent with the apparent inability of *T. gallinae* to phagocytose *E. coli* cells as observed by TEM. However, both species do come into close contact during co-culture and *E. coli* cells were observed tightly integrated into pockets generated by apparent invagination of the plasma membrane of *T. gallinae* (**Figure 2**). These pocket interactions appear to resemble the early stages of phagocytosis seen in TEM and scanning electron microscopy (SEM) for *T. vaginalis* against fungi (Pereira-Neves & Benchimol, 2007). However, the observed pockets in this work appear to never fully ‘close’ around the *E. coli* cells, implying no complete fusion of distal membrane protrusions of *T. gallinae*. Regardless, such tight interactions suggest specific receptor-dependent cell-cell interactions. Candidate receptors which are involved in these interactions could be identified from the RNAseq differential expression results (e.g. BspA-like, which have been suggested to mediate *Trichomonas*-bacteria interactions (Handrich *et al*., 2019)). Without observing phagocytosis, the pocket interactions beg the question as to why they occur at all. They could represent failed efforts at phagocytosis whereby the best effort is made by *T. gallinae* to engulf the *E. coli* and then the bacteria can evade full engulfment, which has been described for some pathogenic *E. coli* strains targeted by human macrophage (Kaper *et al*., 2004). It is worth noting that the DH5α strain used in these co-cultures does not possess the virulence factors enabling such evasion found amongst the pathogenic *E. coli* (Kaper *et al*., 2004). Alternatively, these pockets could provide opportunities for *T. gallinae* cells to control the immediate extracellular environment at the interface between the two cells and perhaps use secretory or membrane bound enzymes or exosomes to actively interact with the bacterial cells (Twu *et al*., 2013) and extract nutrients from the *E. coli* located in the pocket.

The cell tracking data, provides compelling evidence that there is a specific response from *T. gallinae* cells to the presence of the bacteria. The *T. gallinae* cells in co-culture move significantly (p-value < 0.0001) faster on average throughout the culture compared with the monoculture control (**Figure 1C**). Moreover, all five of *T. gallinae*’s transcripts from the annotated kinesin-II genes and some tubulin-tyrosine ligase-like genes, which are likely involved in flagella motility (Pan & Snell, 2002; Kubo & Oda, 2017), are significantly (p-value < 0.05) upregulated during co-culture with *E. coli* (**Figure 4**).

Cell movement for eukaryotes is seldom random and often has a fundamental purpose in terms of cell function to promote vitality and survival (Ginger *et al*., 2008). The observed increase in motility for *T. gallinae* whilst in co-culture could imply a chemotaxis-like response to the presence of bacteria, whereby the increased motility could either provide *T. gallinae* cells with increased opportunities for interactions with the bacteria, or to avoid them. The observation of the formation of pocket interactions and prolonged cell densities of *T. gallinae* in co-culture with *E. coli* is consistent with the former. Chemotaxis has been observed for *T. vaginalis* (Sugarman & Mummaw, 1988). It will be interesting to further explore our developed live cell tracking approach among a broader range of *Trichomonas*-bacteria interactions in co-cultures with different species of parasites and bacteria.

Collectively the observations from live imaging of the co-cultures do imply physiological interactions between *T. gallinae* and *E. coli*. To gain insights into the molecular basis of these interactions, we used a comparative transcriptomics approach (RNAseq) contrasting the transcriptome of *T. gallinae* in monoculture with the *E. coli* co-culture condition for the 24 hour time point with triplicates (**Figure 1A**) (Wang *et al*., 2009). In total, an average of ≈185,000,000 reads per sample amongst the *T. gallinae* vs *E. coli* co-cultures and controls indicates the presence of sufficient reads for all downstream analyses, including differential expression (Liu *et al*., 2013). The common BCV for the data was low (< 0.3), indicating that the difference observed in the differential expression analysis is likely due to biological differences and not just technical noise (McCarthy *et al*., 2012). Here we focused on investigating *T. gallinae* protein-coding genes characterised by significant modulation in response to the presence of the co-cultured bacteria. Significantly upregulated genes in the differential expression analysis represent genes coding for enzymes and other proteins potentially involved in interactions between the two co-cultured organisms. They could also represent genes involved in stress response pathways activated due to acute stresses induced by the co-culture environment, such as nutrient stress due to the presence of a competitor (de Nadal *et al*., 2011).

### The Antibacterial Toolkit

The NlpC/P60 enzymes from the *Trichomonas* genus are a good case study to use to initially query the RNAseq for *T. gallinae*-bacterial interactions as they have been demonstrated to be bacterial targeting enzymes, cleaving the peptide cross links in the PG cell wall of *E. coli* and *L. gasseri* (Pinheiro *et al*., 2018; Barnett *et al*., 2023). Indeed, 2 of the 9 *T. gallinae* NlpC/P60 enzymes, homologous to the enzymes found in *T. vaginalis*, are significantly upregulated **(Figure 2)**, one from each subclass of *Trichomonas* NlpC/P60 (Class A and Class B)(Barnett *et al*., 2023). Homologues of these can also be found in the *T. tenax* genome, meaning they are a conserved group of bacterial targeting enzymes the *Trichomonas* genus has at its disposal (Pinheiro *et al*., 2018; Barnett *et al*., 2023; Mpeyako *et al*., 2024). It is likely the NlpC/P60 endopeptidases cannot act alone in targeting the PG cell wall and instead more likely works in concert with other PG hydrolases and AMPs to access (to cross the outer membrane of a Gram-negative bacteria) and fully (or more extensively) degrade PGs (!!! INVALID CITATION !!! (Pinheiro et al., 2018; Barnett et al., 2023)). Alongside endopeptidases, three enzyme classes are known to hydrolase bonds within the glycan strand of PG; N-acetylglucosaminidase, lysozyme and lytic transglycosylase (Vollmer *et al*., 2008). Lysozyme and lytic transglycosylase both come under the collective term muramidase and they are active on the same β-1-4 glycosidic bond between Glc*N*Ac and MurNAc in PG (Vollmer *et al*., 2008). Two potential glycoside hydrolase families with lysozyme activity are annotated in the *T. gallinae* genome those being GH25 and GH19. Both families are conserved in *T. vaginalis and T. tenax* (Mpeyako *et al*., 2024) and have all their transcripts upregulated in the *T. gallinae* vs *E. coli* co-culture, with 6 out of 9 of these being significantly upregulated (**Figure 5**). Moreover *T. gallinae* has four NagZ-like GH3s (**Figure 9**), also conserved in *T. vaginalis and T. tenax* (Mpeyako *et al*., 2024), which, if the NagZ prediction is accurate, would have N-acetylglucosaminidase activity enabling cleavage of the other β-1-4 glycosidic bond in PG between MurNAc and Glc*N*Ac (Vollmer *et al*., 2008). The expression of one of these enzymes is also significantly upregulated in *T. gallinae* when in co-culture with *E. coli* (**Figure 5**). Lastly, our analyses of the *T. gallinae* vs *E. coli* RNAseq and differential expression analysis highlighted a significantly upregulated gene encoding a putative AMP for *T. gallinae* (**Figure 5**). It was one of two homologues of the bacteriocin AMP-Lcn972 (Turner *et al*., 2013) (**Figure 5**). Based on sequence comparisons, these likely block the action of the bacterial cell wall precursor molecule lipid II (Martínez *et al*., 2008). Integrating the comparative transcriptomics and genomics analyses implicates five different enzyme/peptide groups: NlpC/P60, GH19, GH25, GH3 and AMP-Lcn972, conserved across *T. gallinae*, *T. tenax* and *T. vaginalis,* potentially mediating interactions between these *Trichomonas* species and bacteria. These enzymes and AMP-Lcn972 are all active on, or predicted to act on, the PG cell wall of bacteria or its precursor Lipid II. The RNAseq data in combination with sequence comparisons suggest that *T. gallinae*, and by evident conservation across the genus *Trichomonas*, employs a range of hydrolytic enzymes that could act in concert to either deconstruct the bacterial cell wall or inhibit their production.

The GH19s were chosen for further characterisation due to high level of expression and the potential for the GH19 enzymes to be lysozymes or chitinases or both (Hoell *et al*., 2006; Wohlkönig *et al*., 2010; Lim *et al*., 2012), meaning these could also be potentially implicated in *Trichomonas–*fungi interactions as well. The *T. gallinae* GH19s are conserved throughout *Trichomonas* and more distantly related Metamonads and appear to be the result of prokaryote to eukaryote LGT (**Figure 7**). They appear in domain analysis as two structural classes in *Trichomonas*, a relatively shorter class A and the relatively longer class B which also contains two tandems potentially PG binding SH3b domains of prokaryotic origin (Mitkowski *et al*., 2019) towards the N-terminus (**Figure 6**). These SH3b domains are also present amongst some of the *T. vaginalis* NlpC/P60 enzymes (Pinheiro *et al*., 2018; Barnett *et al*., 2023). Consistent with the GH19 targeting an extracellular substrate (bacterial PG), *T. gallinae* GH19s either have a predicted SP or SP-like sequences in a GH19 alignment (**Figure S5**) and possess the necessary two glutamate catalytic residues required for GH19 function (**Figure 6**) (Hart *et al*., 1995; Hahn *et al*., 2000). Two class B *Trichomonas* GH19s and a catalytic site mutant for TGA_001274700.1 were successfully cloned, expressed, and purified as recombinants from *E. coli* and subject to downstream functional characterisation. The two tested WT GH19s exhibited lysozyme activity (**Figure S7**) in a crude assay using *Micrococcus luteus* cells as a substrate (Shugar, 1952), yet the mutant did not. However, no function was observed for any of the recombinant proteins in a chitinase assay (**Figure S9**). More robust functional characterisation in the form of *E. coli* PG digestion (Glauner, 1988) and tandem HPLC and mass spectrometry confirms the two tested WT *Trichomonas* GH19s were indeed muramidase, cleaving the β1,4-glycosidic bond between the N-Acetylmuramic acid (MurNAc) and N-Acetylglucosamine (Glc*N*Ac) residues in PG (**Figure 8** and **Figure S8**). The mutant was also confirmed to be an inactive enzyme. Therefore, alongside NlpC/P60, the laterally acquired GH19 class B enzymes can be added to a *Trichomonas’* molecular toolkit to target bacterial cell wall polymers. Detailed sequence comparisons that included the class A *Trichomonas* GH19 also suggest these are active enzymes. These notably lack the SH3b domains, which could be important in binding PG prior to hydrolysis (Mitkowski *et al*., 2019). Without the SH3b domains, they may not be able to hydrolyse PG and could possibly be active against chitin.

*Trichomonas* GH3s were also considered of interest due to potential β-N-acetylglucosaminidase activity which is also involved in the breakdown of the sugar backbone of PG (Vollmer *et al*., 2008). *T. gallinae* is predicted to encode six GH3s (**Figure 9**). There are also homologues of these enzymes encoded by *T. tenax and T. vaginalis* (Mpeyako *et al*., 2024). Multiple sequence alignment and phylogenetic analysis split the *Trichomonas* GH3s into those that are homologous to NagZ and those which appear in an outgroup (**Figure 9**). Similarly to the GH19s, the *Trichomonas* NagZ homologues appear to cluster with bacterial sequences, suggesting these genes are the result of a distinct prokaryote to eukaryote LGT event. This acquisition, unlike GH19, does not appear to be conserved amongst wider metamonads. The NagZ homologues are all similar in their domain organisation (**Figure 10**) and they all contain the NagZ catalytic consensus motif required for function (Bacik *et al*., 2012). It is likely that, similar to other bacterial NagZs, the *Trichomonas* NagZs are only active on partially digested PG fragments, after the hydrolysis by a lysozyme (Cheng *et al*., 2000). Based on these considerations, it is relevant to consider the result of hypothetical combined muramidase (GH19), endopeptidase (NlpC/P60) and NagZ activities; leading to the degradation of PG into monosaccharides (Vollmer *et al*., 2008). One of the predicted breakdown products is free monomeric Glc*N*Ac (Vollmer *et al*., 2008). Notably, this sugar can be used to promote *T. gallinae growth* (**Figure 11A**). Consistent with this growth phenotype, *T. gallinae* expresses enzymes for each step in the metabolic pathway enabling Glc*N*Ac to be committed to glycolysis (**Figure 11C**). Hence, there is good reason for *Trichomonas* to want to consume free Glc*N*Ac as they can use it to make ATP (**Figure 11B**) and promote parasite growth more generally. Based on these considerations, we suggest that the *Trichomonas* GH3 NagZ homologues can be added to the toolkit alongside GH19 and NlpC/P60 to break down bacteria’s PG whilst awaiting functional characterisation.

Notably, our work also identified the first case of a laterally acquired bacterial gene encoding an AMP, which is conserved among *Trichomonas* and some additional Metamonada. The initial identification for these occurred by identifying significantly upregulated genes in the *T. gallinae* vs *E. coli* co-culture, leading to the observation of a log_2_FC of 2.6 for TGA_001351900.1 (p-value < 1.06E-10). This gene was annotated as “Bacteriocin (Lactococcin_972), putative” from the *T. gallinae* A1 annotation. This discovery warranted further bioinformatic investigations. Each *Trichomonas spp.* encodes for two paralogous AMP-Lcn972 that have significant similarity to the AMP-Lcn972 from *Lactococcus spp.* and other bacterial homologues in BlastP searches and that share the Lcn972 domain in profile searches (**Figure 12**). Moreover, a range of closely related (*Tritrichomonas foetus*) and more distantly related metamonads also encode in their genomes related sequences, suggesting a deep LGT event and subsequent conservation within Metamonada followed by intra-species duplications. The *Trichomonas* AMP-Lcn972s are short peptides predicted to form an Ig-like fold (Bork *et al*., 1994) composed of several anti-parallel beta sheets. These predicted structures align well, by MatchMaker (Meng *et al*., 2006), with Lcn972 from *Lactococcus cremoris* (Turner *et al*., 2013), despite some *Trichomonas* AMP-Lcn972 having N-terminal extensions not seen in the *Lactococcus* AMP-Lcn972 (**Figure 14**). These N-terminal extensions are likely to contain SP (**Figure S11** and **Figure 14**). AMP-Lcn972 from *Lactococcus spp.* is known to interact with the bacterial cell wall precursor molecule lipid II, inhibiting the ability to build and remodel the PG cell wall in other closely related lactic acid bacteria species (Martínez *et al*., 1996; Martínez *et al*., 2008). Based on the bioinformatics, the function of the *Lactococcus* Lcn972s and the observation of a significant upregulation of the expression of one AMP-Lcn972 from *T. gallinae* in co-culture with *E. coli* the *Trichomonas* AMP-Lcn972 represents a good target for future functional characterisation. The AMP-Lcn972 represent another molecule originating from LGT, identified by comparative transcriptomics in *T. gallinae* vs *E. coli* co-culture and that could be secreted and target bacteria, through inhibiting cell wall construction and remodelling by interaction with lipid II.

Overall, the combined action of the laterally acquired NlpC/P60, GH19, GH3 and AMP-Lcn972 proteins to hydrolyse the PG wall of bacteria and inhibit their synthesis, represents a conserved molecular toolkit which *Trichomonas* could use to exploit and control bacterial members of the mucosal microbiota. The toolkit could actively contribute to the targeting of bacterial cell wall integrity and contribute to both lysis and/or the release of nutrients for the *Trichomonas* to exploit, including PG-derived monosaccharides, membrane lipids and bacterial intracellular contents upon lysis of the targeted cells.

The *T. gallinae* -*E. coli* pocket interactions identified by TEM (**Figure 2**), represent an intriguing interface in which *T. gallinae* could use the toolkit to effectively ‘attack’ the *E. coli* cells. Integrating our various data supports a model of a bacterial cell wall degradation and disruption hypothesis at the site of the pocket interactions by *Trichomonas*. The use of the toolkit as described here could have implications on microbiota homeostasis and pathogenesis of *Trichomonas*. If these tools have activity on a broad range of bacteria or some biomass-dominant members of the mucosal microbiota, they could contribute to dysbiosis by the targeting of members of the microbiota such as the important *Lactobacillus* mutualistic species of the human urogenital tract, which are known to be effectively phagocytosed by *T. vaginalis* (Rendón-Maldonado *et al*., 1998). In the context of phagocytosed cells, we predict that some members of the molecular toolkit will be targeting bacterial cell walls within the phagolysosomes. Furthermore, during dysbiosis (e.g. BV) the toolkit could contribute to the maintenance of the dysbiotic state by the prevention of the mutualistic microbiota re-emerging/establishing itself. Dysbiosis is a hallmark of disease during infections of *T. gallinae*, *T. vaginalis* and *T. tenax* and the conservation of the toolkit across all three species is suggestive that the *Trichomonas* species could contribute directly to the establishment and/or maintenance/worsening of the dysbiotic phenotype (Brotman *et al*., 2012; Maritz *et al*., 2014; Bisson *et al*., 2019; Ji *et al*., 2020). Any action of the combined toolkit on the microbiota will likely contribute to the release of PG degradation products. Notably, PG degradation products are known to stimulate strong inflammatory responses from the host which in turn can lead to, maintain or worsen dysbiosis and by doing so could be an important factor contributing to the damaging of mucosal surfaces through excessive and chronic inflammations (Humann & Lenz, 2009; Wolf, 2023; Zhao *et al*., 2023).

### Mechanism for Zoonosis

*Trichomonas*, and wider Trichomonads, are regarded as having multiple zoonotic origins and importantly, have the capability for future zoonotic transfer events (Maritz *et al*., 2014). Yet the full mechanism for potential zoonosis amongst trichomonads is not understood. The close clustering of proteins expressed in both humans and avian *Trichomonas* in phylogenetic analyses and the broader phylogeny of *Trichomonas*-like species (Peters *et al*., 2020) strongly suggests both the LGTs of genes encoding the toolkit in a pigeon infecting *Trichomonas* ancestor and a potential route of zoonosis from birds to mammals, including humans (Malik *et al*., 2011; Peters *et al*., 2020). In this study and the related ones on NlpC/P60 endopeptidases (Pinheiro *et al*., 2018; Barnett *et al*., 2023), molecular phylogenies indicate the conservation of the entire hypothetical *Trichomonas* antibacterial toolkit across *Trichomonas*, and for some genes also wider Trichomonads (**Figure 7**, **Figure 9** and **Figure 13**). Moreover, Trichomonads found in clinical samples are not restricted to specific body sites, but each site they have been identified in all share the presence of a microbiota (Maritz *et al*., 2014). It can therefore be hypothesised that *Trichomonas* and wider Trichomonads can utilise their antibacterial toolkit to target and exploit the resident mucosal microbiotas of various hosts and sites. This in turn could facilitate host transfers, including zoonotic transmission events, relying not only on the general environment of the host mucosa but perhaps, key to Trichomonads’ host and mucosal site flexibility, the members of the microbiota itself. In addition to modulating the virulence of *T. vaginalis* against human cells in *in vitro* assays (Margarita *et al*., 2022), it was also suggested that *T. vaginalis* symbioses with mycoplasma-like *M. hominis* and *candidatus* M. girerdii might modulate how the parasite impact the vaginal microbiota (Margarita *et al*., 2020). Consistent with this hypothesis (i) a number of dually infected *T. vaginalis-ca.* M. girerdii women were characterised by a vagina microbiota with low 16SrRNA read counts for other bacteria then *ca.* M. girerdi (Fettweis et al., 2014) and (ii) several genes encoding homologues of the molecular toolkit we identified in this study were significantly up-regulated in *T. vaginalis* harbouring either or both mycoplasma-like bacteria (Margarita *et al*., 2022) (**Figure S14**).

## Conclusion

Our work reveals a conserved and multifaceted antibacterial molecular toolkit amongst *Trichomonas spp.*, enabling these symbionts to engage with and manipulate cell wall-containing bacteria. Starting from our *T. gallinae* and *E. coli* co-culture model we combined transcriptomic, comparative genomic, phylogenetic and molecular analysis to identify the toolkit components, including enzymes (NlpC/P60, GH19 and GH3) and antimicrobial peptides (AMP-Lcn972), that likely cooperate to degrade bacterial PG and inhibit its synthesis. Conservation of the toolkit across *Trichomonas spp.*, and occasionally more broadly in Metamonads, implies an evolutionarily shared mechanism to target, exploit and reshape mucosal bacterial communities across diverse hosts and sites.

We also illustrate how the co-culture system increases the fitness of *T. gallinae* by virtue of cell density persistence and enhanced motility whilst also forming distinct pocket interactions with the *E. coli*. These phenotypic responses coincide with dramatic transcriptomic shifts and upregulation of bacterial targeting effectors. Together, these findings support a model in which *Trichomonas* actively senses and responds to bacterial cues, enabling precise interactions with, and killing of, bacterial cells for nutrient acquisition.

Importantly, in the context of infection, the deployment of the toolkit may disrupt microbiota homeostasis by degrading biomass-dominant or mutualistic bacterial taxa such as *Lactobacillus* species. Such disruptions could contribute to the onset or maintenance (or the worsening) of dysbiosis, including through the release of PG breakdown products known to provoke strong inflammatory responses. This, in turn, may drive chronic damaging mucosal inflammations, hallmarks of trichomonad-associated diseases.

The presence of this toolkit in both avian and human-infecting Trichomonads, and its likely origin via LGTs, raises the possibility that microbiota exploitation could facilitate host switching and zoonotic transmission. Therefore, these findings expand our understanding of *Trichomonas* as active modulators of microbial and host environments, with implications for *Trichomonas* pathogenesis mediated through direct and indirect host-microbiota-immune system interactions. Future studies focusing on the regulation, specificity, and host impact of these effectors will be essential to clarify their roles in infection, inflammation, and interspecies transmission alongside observation of the bacterial response to challenge by *Trichomonas spp.* and their effectors.

## Materials and Methods

### Trichomonas culture

All *Trichomonas* spp. cultures and co-cultures were maintained in 15 mL or 50 mL Falcon tubes sealed with Parafilm, using TYM medium (Clark & Diamond, 2002), and incubated at 37°C in a stationary incubator.

### *E. coli* Culture for recombinant protein expression

Liquid cultures of *E. coli* (DH5α, BL21 DE3 and BL21 DE3 STAR) were grown in Lennox LB medium (Formedium, LBX0104) supplemented with 100 µg/mL carbenicillin for co-cultures or 50 µg/mL kanamycin for protein expression. Cultures were incubated at 37°C and 200 rpm in bafled conical flasks, inoculated by picking a single *E. coli* colony from solid medium using a sterile loop. Antibiotic resistance in each strain was conferred by plasmid-encoded genes.

### *T. gallinae* – *E. coli* (DH5α) co-cultures

All co-culture procedures were performed under aseptic conditions using a lateral flow hood, except for initial bacterial handling, which was conducted under a Bunsen burner flame. On the day of co-culture, a *T. gallinae* culture was confirmed to be at high density (>1×10⁶ cells/mL) by haemocytometry. Prior to the day of co-culture, 10 mL of Lennox LB medium (Formedium, LBX0104) supplemented with 100 µg/mL carbenicillin was inoculated with *E. coli* (DH5α), harbouring a plasmid mediating carbenicillin resistance (pSG1154), by picking a single colony from solid medium. This starter culture was incubated overnight at 37°C, then diluted 1:100 into 100 mL of fresh carbenicillin-supplemented Lennox LB (Formedium, LBX0104) in a baffled conical flask. The culture was incubated for no more than 1 h at 37°C and 200 rpm. Bacterial density (CFU/mL) at this stage was estimated by measuring optical density (A600) and referencing a previously established growth curve for the strain.

To initiate co-culture, *T. gallinae* cells were adjusted to a final density of 2.5 × 10⁵ cells/mL by transferring the appropriate volume of high-density culture into pre-warmed (37°C) fresh TYM medium supplemented with 100 µg/mL carbenicillin. Similarly, the bacterial culture was diluted to achieve a 1:10 *E. coli* to *T. gallinae* cell ratio. This setup was prepared in triplicate to generate three biological replicates. Control cultures lacking either *T. gallinae* or *E. coli* were prepared in parallel using the same method. All cultures were incubated at 37°C in Parafilm-sealed Falcon tubes for up to 48 hrs. Throughout the incubation period, *T. gallinae* density was monitored via haemocytometry, while bacterial density was assessed by serial dilution and spread plating to determine colony-forming units (CFUs). After 24 hrs of co-incubation, up to 90% of the remaining volume (≤45 mL) from each *T. gallinae* monoculture and co-culture was harvested by centrifugation at 2000 × *g* for up to 10 min at room temperature using a swinging-bucket centrifuge (Eppendorf 5810 R, rotor A-4-81). Supernatants were discarded, and pellets were washed once in PBS (Sigma-Aldrich, D8537), then resuspended in an equal volume of RNAlater (Invitrogen, AM7020). Pellet samples in RNAlater were stored overnight at 4°C before transfer to −80°C for long-term storage and subsequent RNA extraction and purification.

### Total RNA extraction, storage and purification

For total RNA extraction from *Trichomonas spp.* monocultures and co-cultures (see Figure 1A for corresponding growth curve), a modified protocol based on the TRIzol reagent method (Invitrogen, 15596026) was developed to ensure the recovery of high-quality total RNA. Pelleted cultures, previously stored frozen in RNAlater™ Stabilization Solution (Invitrogen, AM7020), were used for extraction. All reagent containers, equipment, and workspaces were first cleaned with 70% ethanol, followed by RNaseZap™ RNase Decontamination Solution (Invitrogen, AM9782). Samples in RNAlater were resuspended 1:1 in PBS (Sigma-Aldrich, D8537), and 1 mL aliquots were transferred into Eppendorf tubes. Aliquots were centrifuged at 6000 × g for 5 min at 4°C in a benchtop microcentrifuge, and supernatants were discarded. Pellets were transferred to a fume hood (pre-treated with RNaseZap), and lysed by adding 0.5 mL TRIzol reagent (Invitrogen, 15596026), with pipetting to homogenize. Lysates were incubated at room temperature for 5 min to ensure complete lysis. Subsequently, 0.1 mL chloroform (Sigma-Aldrich, 288306) was added, and tubes were inverted by hand for 3 minutes, followed by centrifugation at 16,100 × g for 15 min at 4°C (Eppendorf 5415 R, rotor F45-24-11). After centrifugation, the clear upper aqueous phase was carefully transferred to a fresh tube, avoiding contamination from interphase or organic layers. Downstream processing was carried out on the bench. To the aqueous phase, 0.5 µL of RNase-free glycogen (Thermo Scientific, R0551) and 0.25 mL isopropanol (Sigma-Aldrich, W292907) were added, followed by incubation on ice for 10 min.

Samples were centrifuged at 16,100 × g for 10 min at 4°C. If no visible pellet formed, centrifugation was repeated until one was apparent. The RNA pellet was washed with 0.5 mL of 75% ethanol (VWR Chemicals, 20821.321) and vortexed briefly to mix. Samples in ethanol were stored at −80°C before downstream processing.

Each ethanol–pellet mixture was centrifuged at 12,000 × g for 10 min at 4°C (Eppendorf 5415 R, rotor F45-24-11), and the supernatant was discarded. Pellets were air-dried under a Bunsen burner flame for no more than 5 min. Dried RNA pellets were treated with TURBO DNase (Invitrogen, AM19070), according to the manufacturer’s instructions, to remove residual genomic DNA. Following DNase treatment, RNA-containing supernatants (in TURBO DNase buffer) were pooled by original culture origin into fresh RNase-free tubes. All pooled RNA samples were snap-frozen in liquid nitrogen. An aliquot (5 µL) from each sample was retained for RNA quantification and quality assessment. The remaining RNA was stored at −80°C until submission for RNA sequencing, during which samples were shipped on dry ice.

RNA concentration and purity were assessed using a NanoDrop 2000 spectrophotometer (Thermo Scientific, ND-2000) and a Qubit 4 Fluorometer (Thermo Scientific, Q33238) with the Qubit RNA BR Assay Kit (Invitrogen, Q10210). For RNA sequencing, samples were required to meet a minimum concentration of ≥50 ng/µL in a total volume of ≥20 µL. Purity thresholds were defined as A260/A280 and A260/A230 ratios of approximately 2.0, as measured by NanoDrop.

### RNA sequencing and analysis

Purified total RNA extracted was sent to Novogene for RNAseq sequencing. Ribosomal RNA was first depleted using the Ribo-Zero rRNA Removal Kit (Illumina, 20040526). The remaining RNA underwent library preparation and sequencing on the Illumina NovaSeq 6000 platform (Illumina, 20012850). Library preparation included RNA fragmentation (250–300 bp), reverse transcription to double-stranded cDNA, end repair, poly(A) tailing, adapter ligation, size selection, and PCR amplification. Libraries were quality-checked using a Qubit 4 Fluorometer (Thermo Scientific, Q33238) before sequencing. Post-sequencing, read quality and quantity were initially assessed by Novogene using internal pipelines. The read length was 150 bp. All recovered reads were subsequently re-evaluated in-house to independently verify read quality using the FastQC software (Andrews, 2010). Results were aggregated using MultiQC (Ewels *et al*., 2016) to generate summary files for each RNAseq samples. Once reads were deemed of sufficient quality adapter sequences were removed from the raw read files using the Cutadapt software (Martin, 2011).

### RNAseq data analyses

RNAseq reads from co-culture experimentation of sufficient quality (with adapters removed) were aligned to the *T. gallinae* A1 genome (Alrefaei *et al*., 2019) using the STAR (v2.7.0e) software (Dobin *et al*., 2013) in paired read mode. Both the standard output and the mapped reads per gene table were collected to be used in downstream differential expression analysis. Differential expression analysis was performed using the edgeR package (Robinson *et al*., 2009), in the R environment (R Core Team, 2021), under default parameters. Mapped read count tables were used to compare expression profiles between monoculture and co-culture conditions. Resultant datasets were compiled into individual .tab files, and summary figures were generated within edgeR. Genes with edgeR calculated p-value < 0.05 were considered significantly differentially expressed.

### Cell tracking: Time-lapse Microscopy and Analysis

Time-lapse microscopy was performed using a custom image-capturing protocol and an image analysis pipeline designed specifically for tracking *Trichomonas spp.* cells in co-culture with bacteria. The goal was to exclusively monitor *Trichomonas spp.* cells while distinguishing them from co-cultured bacteria. This image analysis pipeline can be applied to any co-culture system involving *Trichomonas spp.* and a bacterial or fungal species, provided the latter differs significantly in size, shape, motility, and appearance under phase contrast microscopy. The pipeline utilizes only ImageJ (Schindelin *et al*., 2015) and associated plugins or utilities.

Co-cultures of *T. gallinae* and *E. coli* were prepared alongside monoculture controls using the same method described previously. Aliquots (200 µL) of co-culture samples were loaded into 4-chamber culture slides (BD Falcon, 354114) and imaged using a Nikon TiE inverted widefield microscope equipped with a Nikon DS-Fi1 colour camera and a phase contrast lens. For each sample and time point (0 h, 4 h, 24 h, and 48 h), images were captured at regular intervals over a 1.5-minute period, resulting in >1000 images per sample per time point.

Images were converted to 8-bit format and ran through a custom Ilastik (Berg *et al*., 2019) model (generated on randomly selected small regions of the dataset) trained to identify *T. gallinae* cells only, within the culture. Segmentation of output probability maps was optimised by direct comparison to manually annotated example images, resulting in a selected segmentation protocol of pixel value subtraction of 50, Gaussian filtering (sigma=2.0), and globally thresholded according to the “Minimum” algorithm within FIJI. Particles were subsequently excluded if they had an area larger than 40 pixels, or if their circularity was less than 0.25. 3D Gaussian blurring using sigma values of 1.0 in X, Y, and Z axes was then applied to facilitate ease of tracking (Schindelin *et al*., 2015).

Output segmentations were then subjected to tracking using TrackMate (Dmitry *et al*., 2021) to identify paths of individual identified *T. gallinae* cells. The process was run on all available images using through the development of macros for use within FIJI, available at https://github.com/NCL-ImageAnalysis/Molecular-cellular-and-evolutionary-insights-into-Trichomonas-bacteria-Interactions. The output included metrics describing: Track duration (frames), track mean speed (pixels/frame), Track total distance travelled (pixels), Track maximum distance travelled (pixels), Track confinement ratio, mean straight line speed (pixels/frame), linearity of forward progression and mean directional change rate. For final data analysis and presentation, ROUT (Motulsky & Brown, 2006) analysis was used to trim outliners within these large datasets. The trimming step mostly removed extremely high values within the dataset, which were considered to be anomalous.

### Transmission Electron Microscopy

Co-culture samples were prepared for TEM in-house, with final sample processing and imaging performed in collaboration with the Newcastle University Electron Microscopy Research Services, led by Dr Tracy Davey. *T. gallinae–E. coli* co-cultures were centrifuged at 2000 × g for up to 10 min at room temperature until a visible pellet formed. Supernatants were discarded, and cell pellets were fixed in 2.5% glutaraldehyde (Sigma-Aldrich, G7776) in 0.1 M cacodylate buffer. Fixed suspensions were hand-delivered on the same day to the Electron Microscopy Research Services. Upon arrival, supernatants were removed, and pellets were embedded in agarose and processed to form a resin. Samples were post-fixed with 1% osmium tetroxide (OsO₄), sectioned into ultrathin slices, and stored until imaging. TEM images were acquired using a Hitachi HT7800 120 kV microscope equipped with an EMSIS Xarosa CMOS camera.

### Sequence sampling and homologue detection

To identify homologous sequences to *Trichomonas* proteins of interest, including those used in multiple sequence alignments, a combinatory approach was employed. Homology searches were conducted using the NCBI BLAST suite (Sayers *et al*., 2022), both locally and via the NCBI database, as well as through TrichDB (Alvarez-Jarreta *et al*., 2024). Additional searches targeting carbohydrate-active enzymes were performed using the CAZy database (Cantarel *et al*., 2009), accessed manually and via the cazy-webscraper tool (Hobbs *et al*., 2023). Only sequences with BLAST E-values < 0.05 were retained. Query sequences included *Trichomonas* proteins of interest and related homologues identified amongst the above databases.

### Multiple sequence alignment and phylogenetics

All multiple sequence alignments were generated within the Seaview (Gouy *et al*., 2009) tool. Sequences were aligned with either ClustalO (Sievers & Higgins, 2018) or MUSCLE (Edgar, 2004) on a per alignment basis which will be indicated in the legend for a given alignment/ phylogeny figure.

Multiple sequence alignments were initially masked using Gblocks (least stringent settings in Seaview) (Castresana, 2000) to retain conserved regionswith further minimal manual curation to ensure inclusion of well conserved residues. The resulting curated alignments were analysed using the IQ-TREE pipeline (Trifinopoulos *et al*., 2016) under default parameters for phylogenetic reconstruction. Resulting phylogenies were visualized and annotated using the Interactive Tree of Life (iTOL) tool (Letunic & Bork, 2024).

### Alphafold2 Protein Structure Prediction analysis and comparison

All AlphaFold2 protein structural predictions for *Trichomonas* proteins of interest were generated using the ColabFold v1.5.5 server using default parameters (Mirdita *et al*., 2022). Protein structures generated using AlphaFold2 and those of interest extracted from the PDB database (Berman *et al*., 2000) were visualised and annotated using ChimeraX (Meng *et al*., 2006). Protein structure comparison was also conducted within ChimeraX using the MatchMaker tool (Meng *et al*., 2006).

### *Trichomonas* gDNA extraction

Genomic DNA from *T. gallina* (strain A1, isolate XT-1081/07, clone GF1c) *and T. vaginalis* (G3) cultures was extracted using the DNeasy UltraClean Microbial Kit (Qiagen, 12224-50) following the manufacturer’s protocol. Cell lysis and homogenization were performed using a FastPrep FP120 bead beater (Thermo Electron, FP120A-115). All centrifugation steps were carried out using a benchtop swinging bucket centrifuge (Eppendorf 5415D, rotor: F45-24-11). DNA concentration and purity were assessed using a NanoDrop 2000 spectrophotometer (Thermo Scientific, ND-2000), measuring absorbance at 260 nm and the 260/280 nm ratio. Extracted DNA was further validated via agarose gel electrophoresis.

### Commercial cloning of GH3 (NagZ-like) and AMP-Lcn972

Commercial cloning of GH3 and AMP-Lcn972 into pET30a was performed at GenScript. The ORF for protein of interest focused on the mature proteins without the predicted signal peptide. For the AMP-Lcn972 the ORF focused on the conserved segment that include the Lcn972 domain. The truncated insert for each AMP-Lcn972 used is as follows: aa61-136 tAMP1, aa33-101 tAMP2, aa60-137 gAMP1, aa42-119 gAMP2, aa26-103 vAMP1 and aa35-99 vAMP2. The truncated insert for each GH3 used is as follows: aa29-555 TGA_002045800.1 and aa29-553 TVAGG3_0997370. All constructs were confirmed by sequencing (GenScript).

### Cloning into expression vector and site directed mutagenesis of GH19

For cloning, pET28a was linearized by restriction digestion to facilitate the insertion of DNA fragments of interest (NdeI and XhoI, Thermo Scientific, ER0581 and ER0691 respectively). Primers for PCR for *T. gallinae* and *T. vaginalis* GH19 (TGA_001274700.1 and TVAGG3_0677660 and mutagenesis primers for *T. gallinae* GH19 (TGA_001274700.1MUT1 [E202Q]) are found in supplementary table S3. All PCR amplifications of DNA sequences before cloning were performed using CloneAmp HiFi PCR Premix (Takara, 639298), following the manufacturer’s recommendations. PCR reactions were run in a C1000 Touch Thermal Cycler (Bio-Rad, 1851148) using cycling parameters adjusted according to the length of the target sequence and the CloneAmp HiFi PCR Premix protocol. Ligation of insert into the plasmid vector was performed using the In-Fusion® Snap Assembly EcoDry™ kit (Takara, 638954), following the manufacturer’s “In-Fusion Cloning Procedure. Site-directed mutagenesis was performed using the QuikChange II Site-Directed Mutagenesis Kit (Agilent, 200523), following the manufacturer’s protocol. Mutagenesis primers were designed using the manufacturer’s web-based tool. All constructs were confirmed by Sanger sequencing.

### Recombinant protein expression and purification

For the expression and purification of all recombinant His-tagged *Trichomonas* GH19s (TGA_001274700.1, TGA_001274700.1MUT1 [E202Q] and TVAGG3_0677660), the expression strain *E. coli* BL21 DE3 was used. A 10 ml Lennox LB culture supplemented with 50 µg/mL Kanamycin (Formedium, KAN0025) was inoculated with an individual colony of *E. coli* BL21 DE3 transformed with the pET28a plasmid containing the *Trichomonas* GH19 insert. The culture was grown overnight at 37°C, 200 rpm in a shaking incubator. The next day, a suitable volume of fresh Lennox LB with 50 µg/mL Kanamycin (Formedium, KAN0025) was added to baffled conical flasks, inoculated with 1:20 of the overnight culture, and grown at 37°C, 200 rpm, until an OD (A600) of 0.6-0.8 was reached. Induction with 0.1 mM IPTG (Thermo Scientific, R0392) was then performed. After induction, cultures were incubated overnight at 16°C, 200 rpm. Upon completion of the incubation, cultures were centrifuged based on the culture volume:

- For volumes ≤200 ml, centrifugation was done at 5000 x g for 10 minutes at room temperature in a fixed-angle centrifuge (Eppendorf, Centrifuge 5804 R, rotor: F-34-6-38).
- For volumes >200 ml, centrifugation was done at 8000 rpm for 30 minutes in a high-performance fixed-angle centrifuge (Avanti, J-26 XP, rotor: JLA 8.1).

The supernatant was discarded, and the pellet was resuspended in 10 ml of TALON buffer and homogenised by vortexing. The suspension was kept on ice and subjected to ultrasonic lysis for 2 minutes with 0.5-second pulses using a sonicator (B. BRAUN, LabSonic U). The lysate was then centrifuged at 12,000 x g at 4°C for 10 minutes. The supernatant was separated, with 0.5 ml aliquoted as cell-free extract (CFE) and kept on ice. If the supernatant was particularly turbid, it was filtered through a 0.2 µm syringe filter (Starlab, E4780-1226). The pellet was resuspended in 10 ml of TALON buffer and stored on ice. A gravity flow column was pre-loaded with 4 ml of TALON® Metal Affinity Resin (Takara, 635653) and washed three times with TALON buffer. The column was capped and ready for protein purification. The supernatant was carefully applied to the resin, and the flow-through was collected in a tube labelled Flow Through (FT). The resin was then washed twice with 5 ml of TALON buffer, and the flow-through was collected and named WASH (W). Next, the resin was washed with 4 ml of 10 mM Imidazole solution, and the flow-through was collected and named 10 mM. The resin was then washed with 4 ml of 100 mM Imidazole solution, and the flow-through was collected and named 100 mM-1. This was repeated for a second wash, with the flow-through collected and named 100 mM-2. All flow-through samples were kept on ice. After aliquoting and preparing samples for SDS-PAGE, the remaining volumes of each flow-through were stored at 4°C for up to one month. SDS-PAGE (12%) was used throughout this work to validate recombinant protein expression and purification protocols.

Alternative approaches were explored for the expression and purification of recombinant GH3s and AMP-Lacn972s encoded on a pET30a vector. These included variations of the above expression and purification method, such as altering the *E. coli* expression strain to BL21 STAR. Despite these adjustments, none of the tested approaches were successful in generating purified recombinant soluble proteins.

### Recombinant protein concentration

Fore enzymatic assays, recombinant protein samples were concentrated with the use of Amicon® Ultra-15 Centrifugal Ultracel-30 Filter Units (Millipore, UFC903008). The centrifugal filter with recombinant protein solution was centrifuged at 5000xg in a fixed angle centrifuge (Eppendorf, Centrifuge 5804 R, rotor: F-34-6-38) for five minute time intervals to concentrate.

### Lysozyme enzymatic assay

Protein concentration of purified recombinant proteins was quantified using a NanoDrop 2000 spectrophotometer (Thermo Scientific, ND-2000) prior functional assays. Purified recombinant *Trichomonas* GH19s and mutant GH19 (TGA_001274700.1, TVAGG3_0677660, and TGA_001274700.1MUT1) were subjected to a lysozyme enzymatic assay as described by Shugar et al. (Shugar, 1952) and adapted from the protocol available at https://www.sigmaaldrich.com/GB/en/technical-documents/protocol/protein-biology/enzyme-activity-assays/enzymatic-assay-of-lysozyme to determine lysozyme-like activity. The principle of the assay relies on the lysis of *Micrococcus lysodeikticus*, a gram-positive coccus, when exposed to lysozyme. This lysis produces a ΔA450 of -0.001 per minute at pH 6.24 and 25°C in a 2.6 ml reaction mixture. The protocol was slightly modified to enable the assay to be performed in 96-well plates. A suspension of 0.015% (w/v) *Micrococcus lysodeikticus* (Sigma Aldrich, M37770) in lysozyme enzymatic assay buffer was prepared. 190 µl of this suspension was pipetted into wells on a 96-well plate. Then, 10 µl of a given recombinant protein sample, HEWL (Sigma Aldrich, 62970-1G-F) positive control solution, or blank solution (100 mM imidazole buffer) was added to the Micrococcus suspension in triplicate. The OD (A450) for each well was measured over time for 10 minutes in a plate reader at 25°C.

### Chitinase activity assay

Pure recombinant *Trichomonas* GH19s and mutant GH19 (TGA_001274700.1, TVAGG3_0677660, and TGA_001274700.1MUT1) were subjected to the Chitinase activity assay using the Chitinase Assay kit (Sigma Aldrich, CS0980). In principle, the assay involves the cleavage of 4-nitrophenol-tagged chitin substrates by a chitinase enzyme at basic pH. This cleavage releases 4-nitrophenol, which can be measured colourimetrically at 405 nm. The protocol provided with the kit was followed exactly. OD A405 measurements were taken in triplicate for each substrate with each recombinant protein sample, chitinase positive control solution (Chitinase from *Trichoderma viride* [Sigma Aldrich, C6242]), and blank solution (100 mM imidazole buffer).

### Peptidoglycan degradation assay, HPLC and Mass spectrometry

PG was purified and analysed by HPLC as previously published (Glauner, 1988). 10 µM of a given purified recombinant protein of interest and cellosyl (as positive control, gift from Hoechst, Frankfurt, Germany) was incubated ON with PG from *E. coli* BW25113 or MC1061 in 20 mM sodium acetate buffer pH 5.0. The reaction was stopped by heating to 100°C for 10 minutes in a heat block. Reaction mixtures were centrifuged at 14,000 rpm for 10 minutes at room temperature in a benchtop microcentrifuge and the supernatant collected. Samples were reduced by adding the same volume of 0.5 M sodium borate buffer pH 9.0 and a ‘granule’ of sodium borohydride (Sigma Aldrich, 213462) and incubated for 30 min at room temperature. The pH was adjusted to 3-4 and samples were subject to reversed-phase HPLC as published (Glauner, 1988). Major fractions were collected for mass spectrometry (MS) analysis to generate the MS and MS/MS spectra as published (Bui *et al*., 2009).

### Growth Curves

Growth experiments for (i) *T. gallinae* in the presence of different sugars and (ii) *E. coli* BL21* expressing *Trichomonas* AMP-Lnn972 were generated by filling wells of 96-well plates with 200µl of freshly inoculated culture media. Plates would then be incubated at 37°C in a plate reader (Promega, GloMax Discover GM3000) with intermittent agitation. At 30-minute time intervals, the OD (A600) for each well was measured for up to 48 hrs to follow growth.

### Luminescence ATP viability assay

Pre-cultures of *T. gallinae* were generated by pelleting 6×10^6^ cells (quantified by haemocytometry) at 1.4 k x g for 5 minutes at RT, resuspending in 15 ml pre-warmed modified TYM, and incubating at 37°C, sealed in parafilm for 24 hours. To induce starvation stress, *Trichomonas* cells were pelleted at 1400 x g for 5 minutes at room temperature, washed with PBS and suspended in 1 ml HBSS supplemented with 1% (w/v) monosaccharide. The cell density of suspensions was quantified by haemocytometry, and diluted using the same buffer to a density of 2.5×10^5^ cells/ml in a total volume of 1.5 ml, sealed in parafilm and incubated at 37°C. ATP viability assays were performed using the CellTiter-Glo® Luminescent Cell Viability Assay (Promega), according to the manufacturer’s instructions. Luminescence was measured using a Synergy H1 microplate reader (Biotek), using a digital gain of 135, integration time of 1 second and read height of 1 mm. Results were normalised by subtracting luminescence intensity measured from cell-free control assays performed in parallel, in the same buffer.

## Acknowledgements

We thank Dr Kevin Tyler (University of East Anglia) for providing the annotation file for *T. gallinae* draft genome (strain A1, isolate XT-1081/07, clone GF1c). We thank Dr Emma Foster from the bioimaging facility for training and advice, Dr Tracy Davey from the Electron Microscopy Facility for carrying out the TEM, training and advice, Dr Daniela Vollmer for preparing the purified peptidoglycan and Prof. Henrik Strahl for general advice about *Trichomonas* AMP-Lcn972 (all from Newcastle University). This research made use of the Rocket High Performance Computing service at Newcastle University. Funding: AJH and NPB were supported by the UK Biotechnology and Bioscience Research Council Doctoral Training Partnership program for Newcastle, Liverpool and Durham (grant numbers: BB/T008695/1 and BB/M011186/1 respectively, RPH supervisor). LAM was supported by a Commonwealth Scholarship Commission in the UK (grant number CMCS-2019-109, RPH supervisor) and Newcastle University. Work in the WV lab was funded by the UK Biotechnology and Biological Sciences Research Council (grant number: BBSRC, BB/W013630/1).

## Data availability statement

The RNAseq data are available at the NCBI SRA database (Bioproject: PRJNA1019275, accession numbers SAMN37478063 to SAMN37478068). All the key outputs of the sequence analyses for this study can be found in the Supplementary material. They are all described in the main text and in the Supplementary material.

## Notes

### Competing Interest Statement

The authors have declared no competing interest.

