## Supplementary Figure/Tables for "An evolutionarily conserved laterally acquired toolkit enables microbiota targeting by *Trichomonas*"

### Supplementary Figures:

**Figures S1: Growth of *E. coli* in co-culture with *T. gallinae*.**

**Figures S2: Summary of cell tracking data in *T. gallinae* co-cultures.**

**Figures S3: Distance travelled by *T. gallinae* cells in co-culture with *E. coli.***

**Figures S4: RNAseq read quality assessment**.

**Figures S5: N-terminal alignment of *T. gallinae* GH19 proteins and SH3b domain alignment with *T. vaginalis* NlpC/P60*.***

**Figures S6: Broad GH19 phylogeny.**

**Figures S7: *Trichomonas* GH19 Lysozyme enzymatic assay.**

**Figures S8: Tandem liquid chromatography–mass spectrometry of GH19-digested peptidoglycan degradation products**

**Figures S9: *Trichomonas* GH19 Chitinase activity assay.**

**Figures S10: Phylogenetic analysis, multiple sequence alignment and domain architecture of GH3 proteins.**

**Figures S11: Domain architecture and N-terminal alignment of *Trichomonas* AMP_Lcn972s.**

**Figures S12: Induced and non-induced growth curves for BL21 STAR transformed with *Trichomonas* AMP-Lcn972 expression constructs.**

**Figures S13: SDS-PAGE summary of *Trichomonas* AMP-Lcn972 expression in BL21 STAR *E. coli.***

**Figures S14: Differential expression of *T. vaginalis* antibacterial toolkit genes in the presence of *Mycoplasma* endosymbionts.**

### Supplementary tables:

**Table S1: *T. gallinae-E. coli* co-culture RNAseq read summary data.**

**Table S2: BLASTp hit list for *Trichomonas* AMP-Lcn972 protein TGA_001351900.1 against the NCBI nr database.**

**Table S3: PCR and mutagenesis primers for *Trichomonas* GH19 genes.**

**Table S4: Master table *T. gallinae* gene annotations along with all the RNAseq data counts. See separate excel table.**

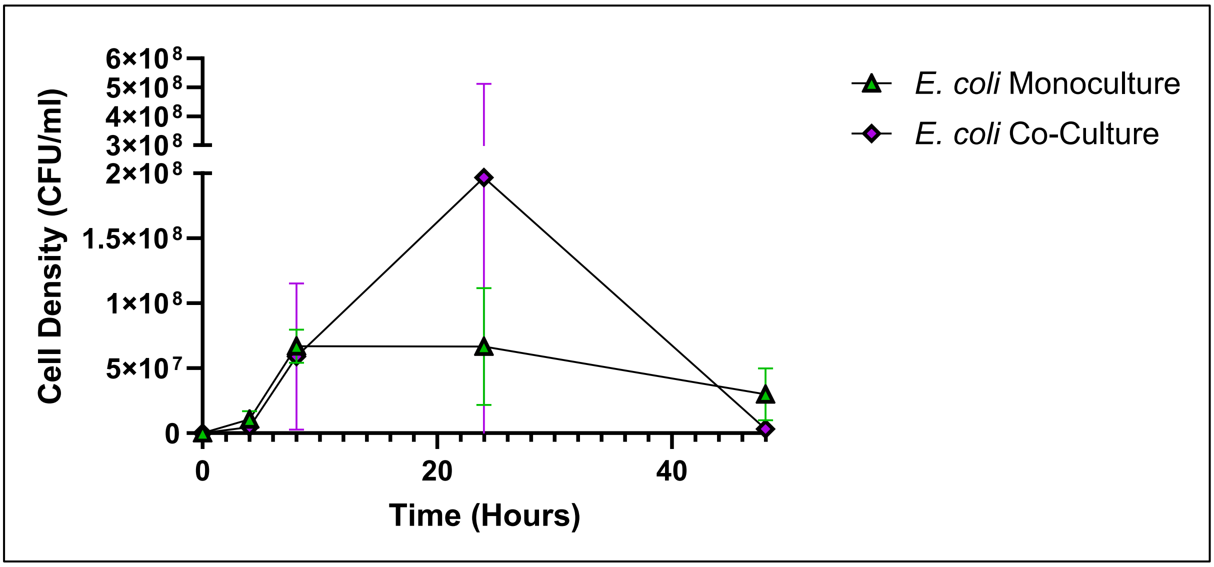


**Figure S1**: **Growth of *E. coli* in co-culture with *T. gallinae*.** Growth curve showing *E. coli* (CFU/ml) in the absence (green) or presence (purple) of *T. gallinae* over 48 hours. Error bars represent standard deviation, n = 3. These data correspond to the same experiment illustrated in Figure 1A (*T. gallinae* cell counts).

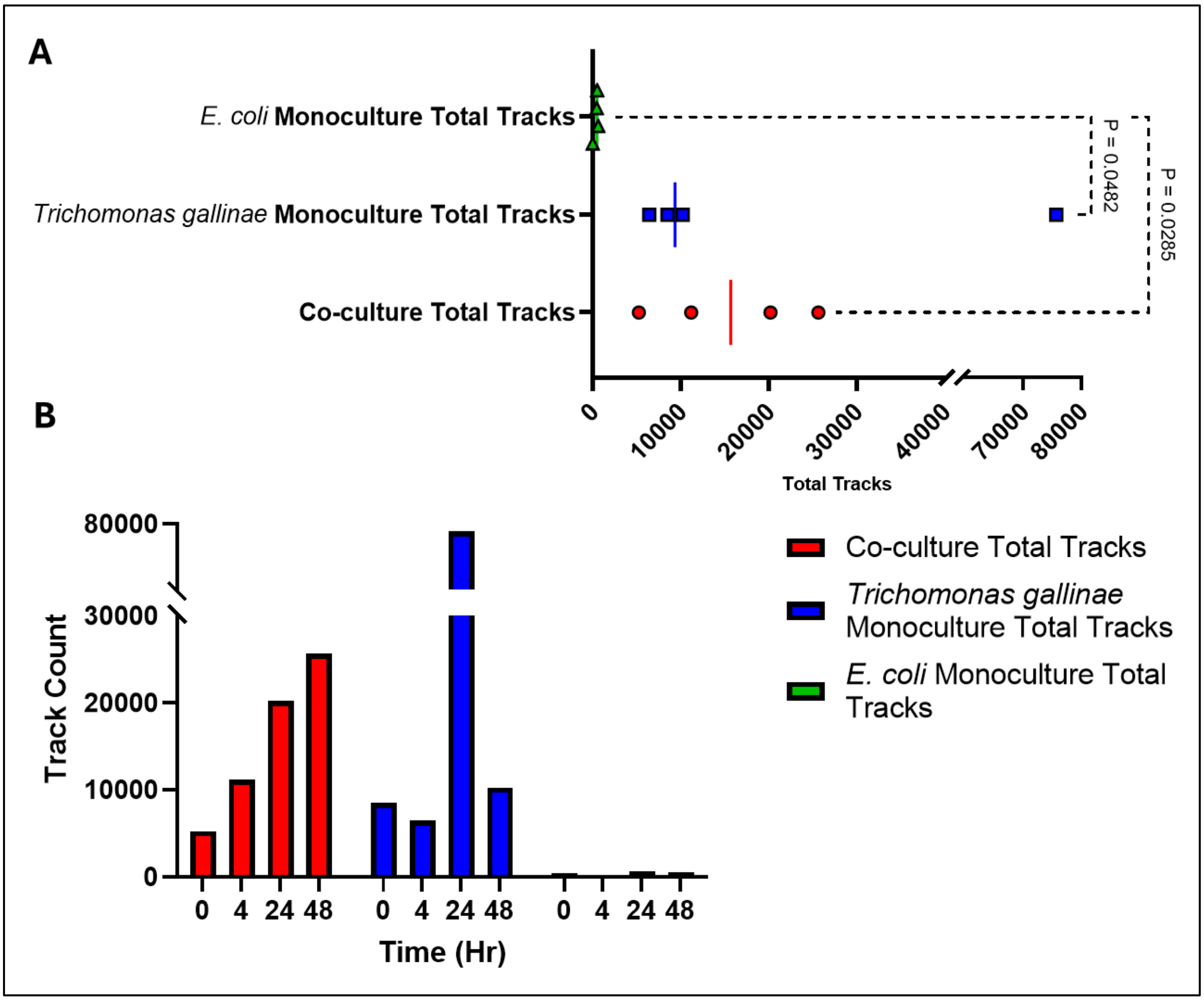


**Figure S2: Summary of cell tracking data in *T. gallinae* co-cultures.** (A) Scatter dot plot showing the total number of tracks detected across four time points for *E. coli* monoculture (green), *T. gallinae* monoculture (blue), and *T. gallinae* – *E. coli* co-culture (red). P-values from Kruskal–Wallis tests are shown to the right of each comparison. Vertical lines indicate median values across all time points for each condition. (B) Bar chart displaying total track count values per time point for *E. coli* monoculture (green), *T. gallinae* monoculture (blue), and co-culture (red).

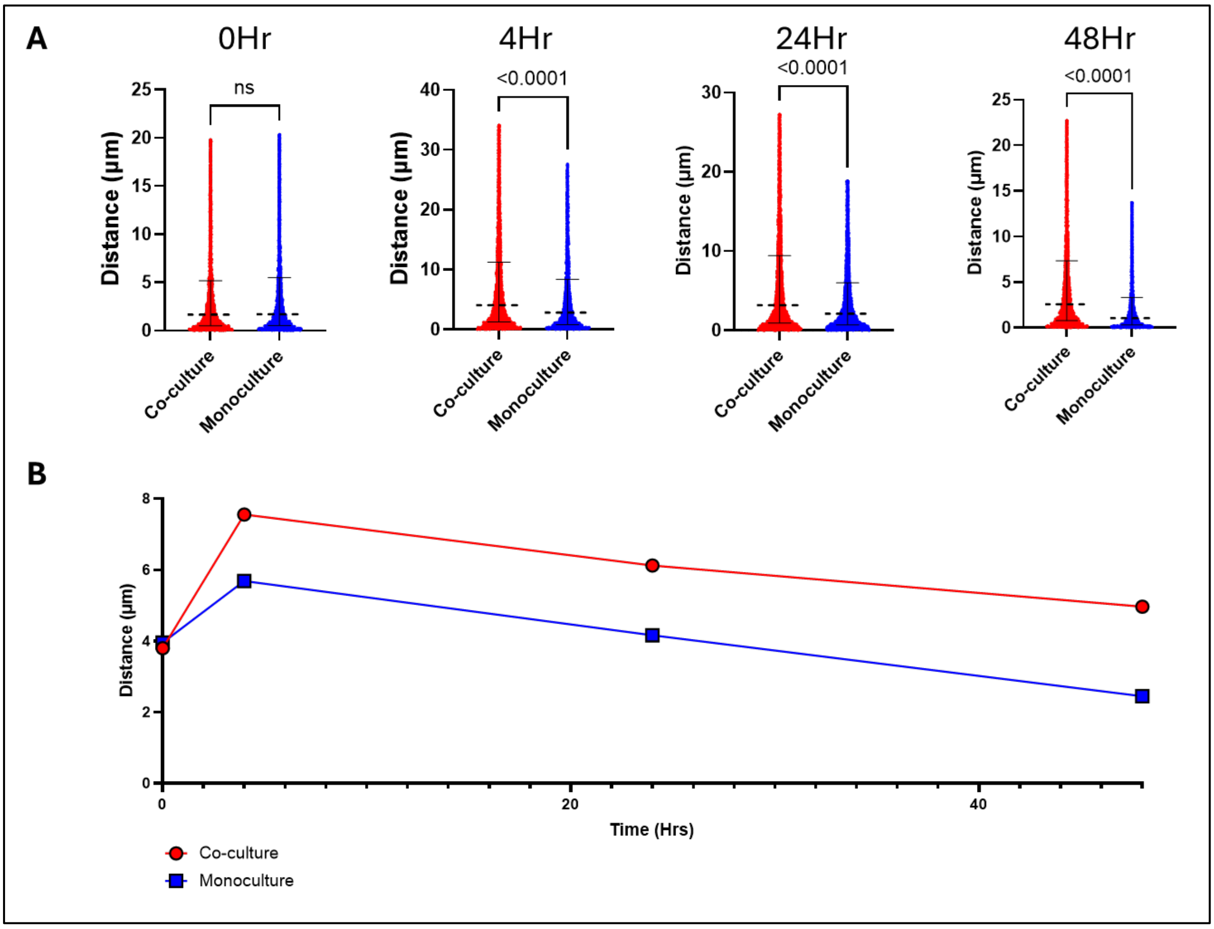


**Figure S3: Distance travelled by *T. gallinae* cells in monoculture or co-culture with *E. coli.*** (A) Violin plots showing *T. gallinae* distance travelled (μm) at multiple time points, measured via live-cell imaging in the absence (blue) or presence (red) of *E. coli*. P-values from Kruskal–Wallis tests are displayed above each comparison. (B) Mean *T. gallinae* distance travelled (μm) over time in monoculture (blue) and co-culture (red) conditions.

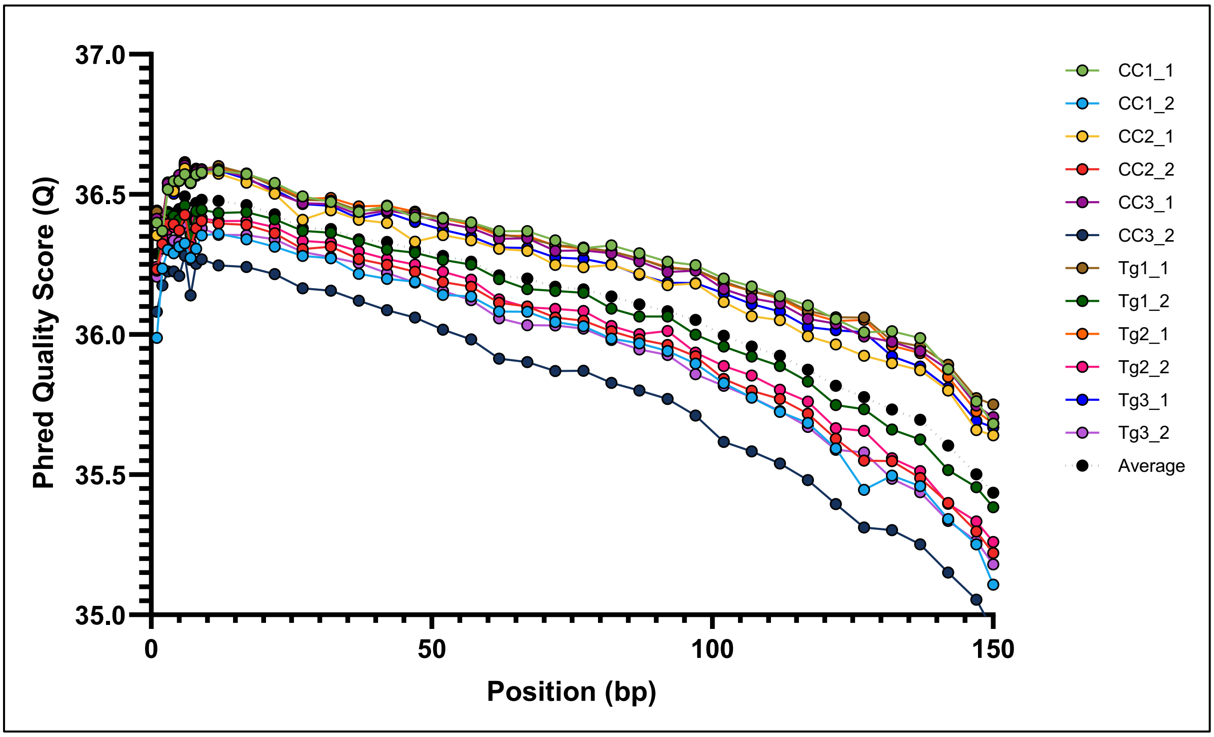


**Figure S4**: **RNAseq read quality assessment**. Average per-base Phred quality scores for each paired-end RNAseq read from *T. gallinae* monoculture and co-culture samples. Co-culture samples are labelled with the prefix "CC" and monoculture samples with the prefix "Tg" both followed by the sample number and either "_1" or "_2" to indicate read pair. Each sample is color-coded individually, with the overall average shown in black.

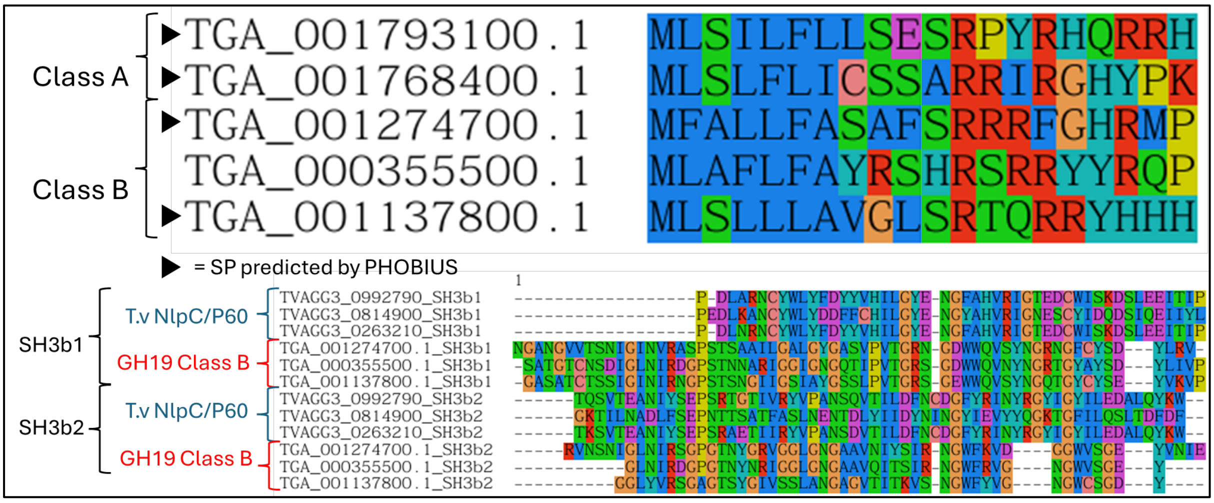


**Figure S5: N-terminal alignment of *T. gallinae* GH19 proteins and SH3b domain alignment with *T. vaginalis* NlpC/P60.** Top: N-terminal alignment of *T. gallinae* GH19 proteins showing conserved sequence features. Black arrows indicate sequences with PHOBIOUS predicted signal peptides (SP). GH19 class designations for each sequence are shown to the left. Bottom: Alignment of SH3b domains from *T. vaginalis* NlpC/P60 proteins and Class B *T. gallinae* GH19 proteins. Each SH3b domain is labelled on the left to indicate its origin.

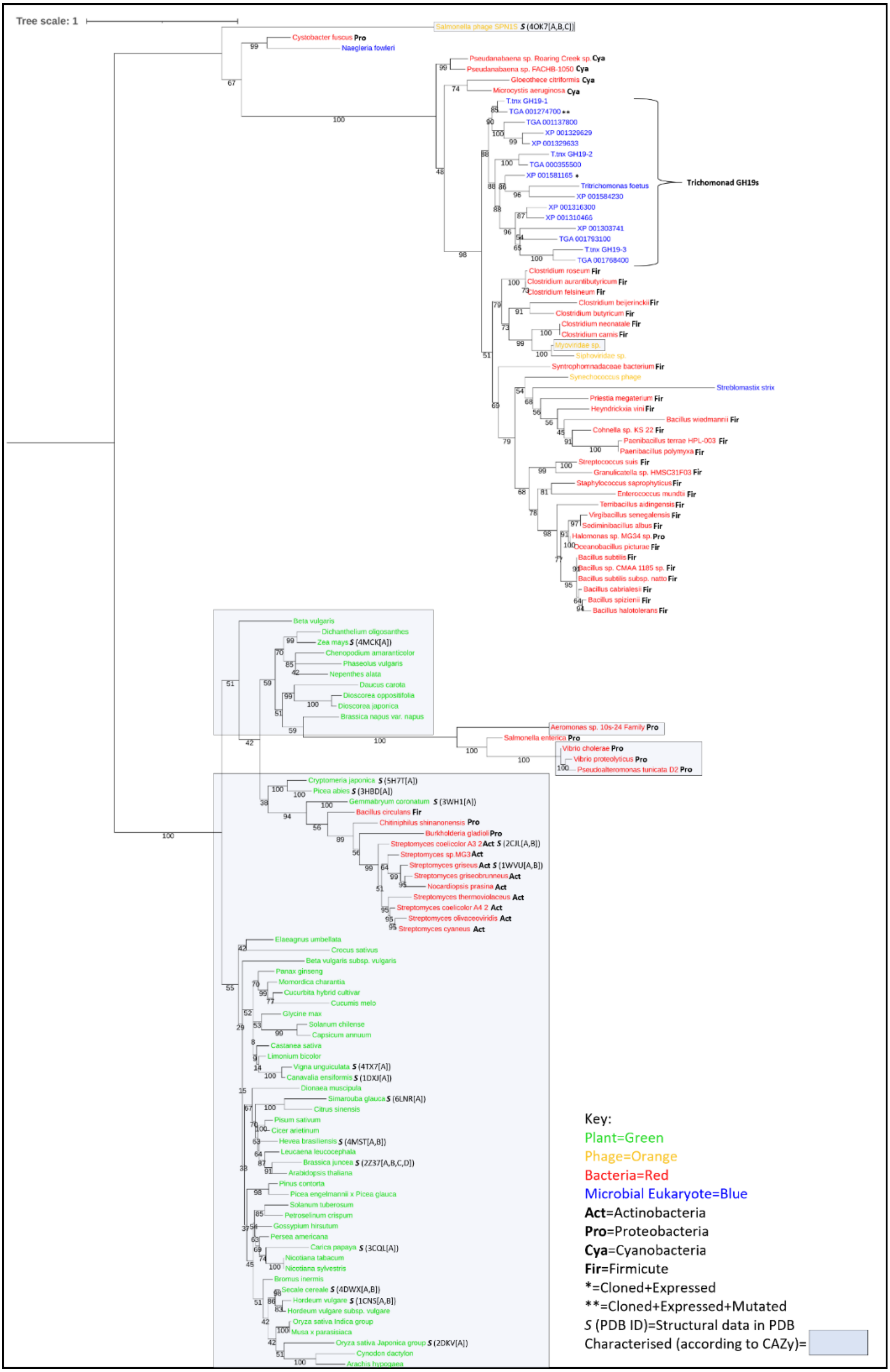

Tree scale: 1
Salmonella phage SPN1S S (4OK7[A,B,C])
Cystobacter fuscus Pro
Naegleria fowleri
Pseudanabaena sp. Roaring Creek sp. Cya
Pseudanabaena sp. FACHB-1050 Cya
Gloeothece citriformis Cya
Microcystis aeruginosa Cya
T.tnx GH19-1
TGA 001274700 **
TGA 001137800
XP 001329629
XP 001329633
T.tnx GH19-2
TGA 000355500
XP 001581165 *
Tritrichomonas foetus
XP 001584230
XP 001316300
XP 001310466
XP 001303741
TGA 001793100
T.tnx GH19-3
TGA 001768400
Trichomonad GH19s
Clostridium roseum Fir
Clostridium aurantibutyricum Fir
Clostridium felsineum Fir
Clostridium beijerinckii Fir
Clostridium butyricum Fir
Clostridium neonatale Fir
Clostridium carnis Fir
Myoviridae sp.
Siphoviridae sp.
Syntrophomonadaceae bacterium Fir
Synechococcus phage
Streblomastix strix
Priestia megaterium Fir
Heyndrickxia vini Fir
Bacillus wiedmannii Fir
Cohnella sp. KS 22 Fir
Paenibacillus terrae HPL-003 Fir
Paenibacillus polymyxa Fir
Streptococcus suis Fir
Granulicatella sp. HMSC31F03 Fir
Staphylococcus saprophyticus Fir
Enterococcus mundtii Fir
Terribacillus aidingensis Fir
Virgibacillus senegalensis Fir
Sediminibacillus albus Fir
Halomonas sp. MG34 sp. Pro
Oceanobacillus picturae Fir
Bacillus subtilis Fir
Bacillus sp. CMAA 1185 sp. Fir
Bacillus subtilis subsp. natto Fir
Bacillus cabrialesii Fir
Bacillus spizizenii Fir
Bacillus halotolerans Fir
Beta vulgaris
Dichanthelium oligosanthes
Zea mays S (4MCK[A])
Chenopodium amaranticolor
Phaseolus vulgaris
Nepenthes alata
Daucus carota
Dioscorea oppositifolia
Dioscorea japonica
Brassica napus var. napus
Aeromonas sp. 10s-24 Family Pro
Salmonella enterica Pro
Vibrio cholerae Pro
Vibrio proteolyticus Pro
Pseudoalteromonas tunicata D2 Pro
Cryptomeria japonica S (5H7T[A])
Picea abies S (3HBD[A])
Gemmabryum coronatum S (3WH1[A])
Bacillus circulans Fir
Chitiniphilus shinanonensis Pro
Burkholderia gladioli Pro
Streptomyces coelicolor A3 2 Act S (2CJL[A,B])
Streptomyces sp. MG3 Act
Streptomyces griseus Act S (1WVU[A,B])
Streptomyces griseobrunneus Act
Nocardiopsis prasina Act
Streptomyces thermoviolaceus Act
Streptomyces coelicolor A4 2 Act
Streptomyces olivaceoviridis Act
Streptomyces cyaneus Act
Elaeagnus umbellata
Crocus sativus
Beta vulgaris subsp. vulgaris
Panax ginseng
Momordica charantia
Cucurbita hybrid cultivar
Cucumis melo
Glycine max
Solanum chilense
Capsicum annuum
Castanea sativa
Limonium bicolor
Vigna unguiculata S (4TX7[A])
Canavalia ensiformis S (1DXJ[A])
Dionaea muscipula
Simarouba glauca S (6LNR[A])
Citrus sinensis
Pisum sativum
Cicer arietinum
Hevea brasiliensis S (4MST[A,B])
Leucaena leucocephala
Brassica juncea S (2Z37[A,B,C,D])
Arabidopsis thaliana
Pinus contorta
Picea engelmannii x Picea glauca
Solanum tuberosum
Petroselinum crispum
Gossypium hirsutum
Persea americana
Carica papaya S (3CQL[A])
Nicotiana tabacum
Nicotiana sylvestris
Bromus inermis
Secale cereale S (4DWX[A,B])
Hordeum vulgare S (1CNS[A,B])
Hordeum vulgare subsp. vulgare
Oryza sativa Indica group
Musa x parasisiaca
Oryza sativa Japonica group S (2DKV[A])
Cynodon dactylon
Arachis hypogaea
Key:
Plant=Green
Phage=Orange
Bacteria=Red
Microbial Eukaryote=Blue
Act=Actinobacteria
Pro=Proteobacteria
Cya=Cyanobacteria
Fir=Firmicute
*=Cloned+Expressed
**=Cloned+Expressed+Mutated
S (PDB ID)=Structural data in PDB
Characterised (according to CAZy)=

**Figure S6:** A **broad GH19 phylogeny.** Midpoint-rooted phylogenetic tree generated from a masked multiple sequence alignment of *Trichomonas* and trichomonad GH19 proteins, along with a broad range of homologues identified via BLASTp searches against the NCBI's nr database and characterised GH19S from the CAZy database. The maximum likelihood phylogeny was inferred using the LG+I+G4 model with ultrafast bootstrapping; bootstrap values >70 are displayed at their respective nodes. The scale bar shows the inferred number of substitutions per site. Branch labels indicate the species of origin and either a PDB ID (if structural data is available) or the *Trichomonas* gene ID. Tree annotations follow the key provided in the bottom right corner.

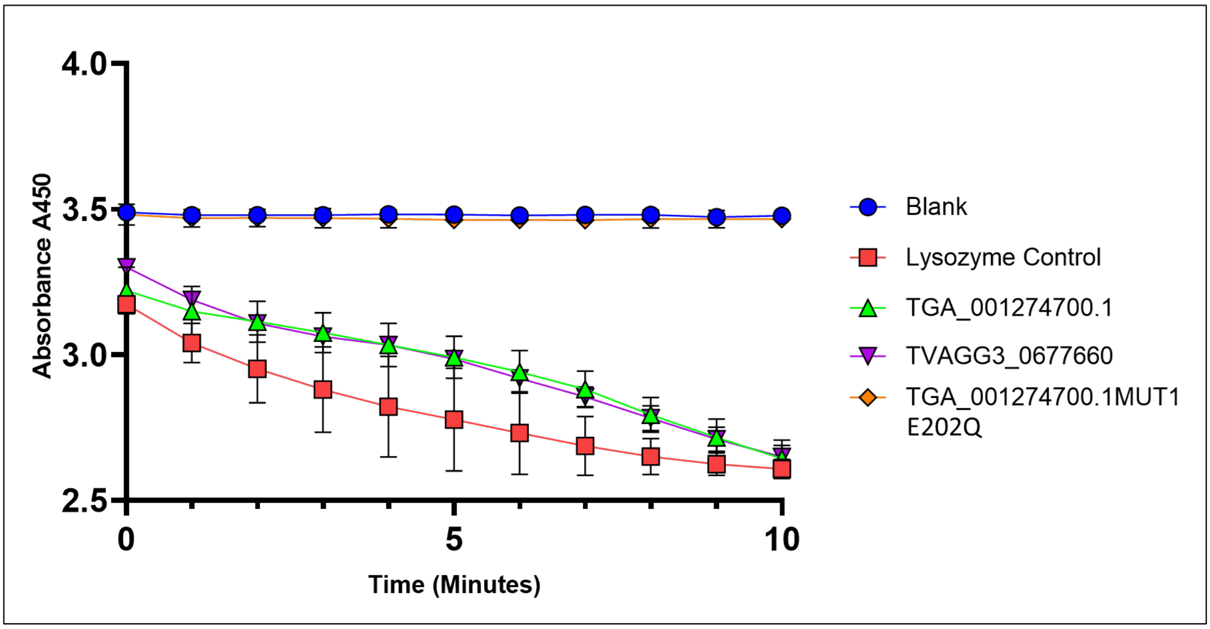


**Figure S7:** ***Trichomonas* GH19 Lysozyme enzymatic assay.** OD (A450) values over time for a lysozyme activity assay using *Micrococcus luteus* as the substrate. Conditions include: lysozyme control (red, hen egg white lysozyme, GH22), recombinant *T. gallinae* GH19 (TGA_001274700.1, green), recombinant *T. vaginalis* GH19 (TVAGG3_0677660, purple), catalytic mutant TGA_001274700.1MUT1-E202Q (orange), and a negative control (Blank, blue). N = 8 and error bars represent standard deviations.

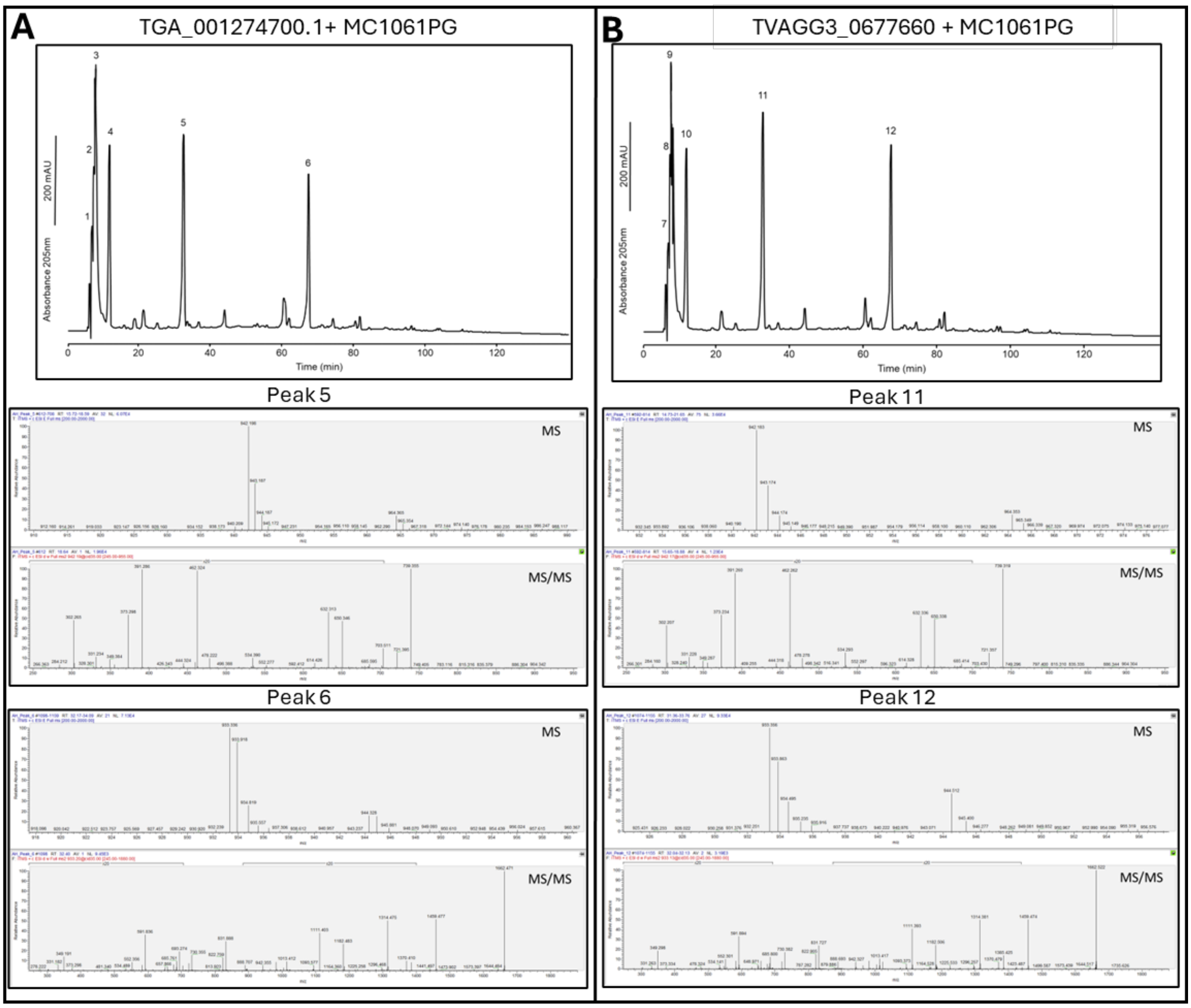


**Figure S8: Tandem liquid chromatography–mass spectrometry of GH19-digested peptidoglycan degradation products.** HPLC chromatograms displaying degradation products generated from *E. coli* MC1061PG peptidoglycan incubated in 20 mM sodium acetate buffer (pH 5.0) with recombinant GH19 proteins TGA_001274700.1 and TVAGG3_0677660. Corresponding tandem MS and MS/MS spectra are shown below each chromatogram. Peaks in the chromatograms are labelled and matched to the appropriate spectra below.

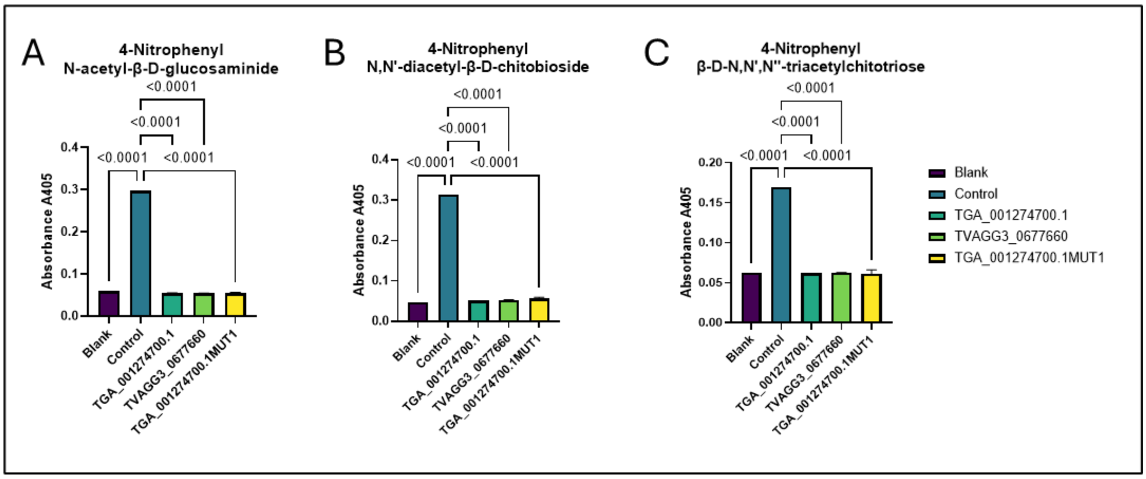


**Figure S9: *Trichomonas* GH19 chitinase activity assay.** Absorbance measurements (A405) showing enzymatic activity on three chitin-based substrates: (A) 4-Nitrophenyl N-acetyl-β-D-glucosaminide, (B) 4-Nitrophenyl N,N'-diacetyl-β-D-chitobioside, and (C) 4-Nitrophenyl β-D-N,N',N''-triacetylchitotriose. Reactions were carried out using a chitinase control (chitinase from *Trichoderma viride*, teal GH18), TGA_001274700.1 (turquoise), TVAGG3_0677660 (green), TGA_001274700.1 MUT1 E202Q (yellow), and a blank control (purple). P-values from one-way ANOVA tests are shown above each comparison. N = 3 and error bars represent standard deviations.

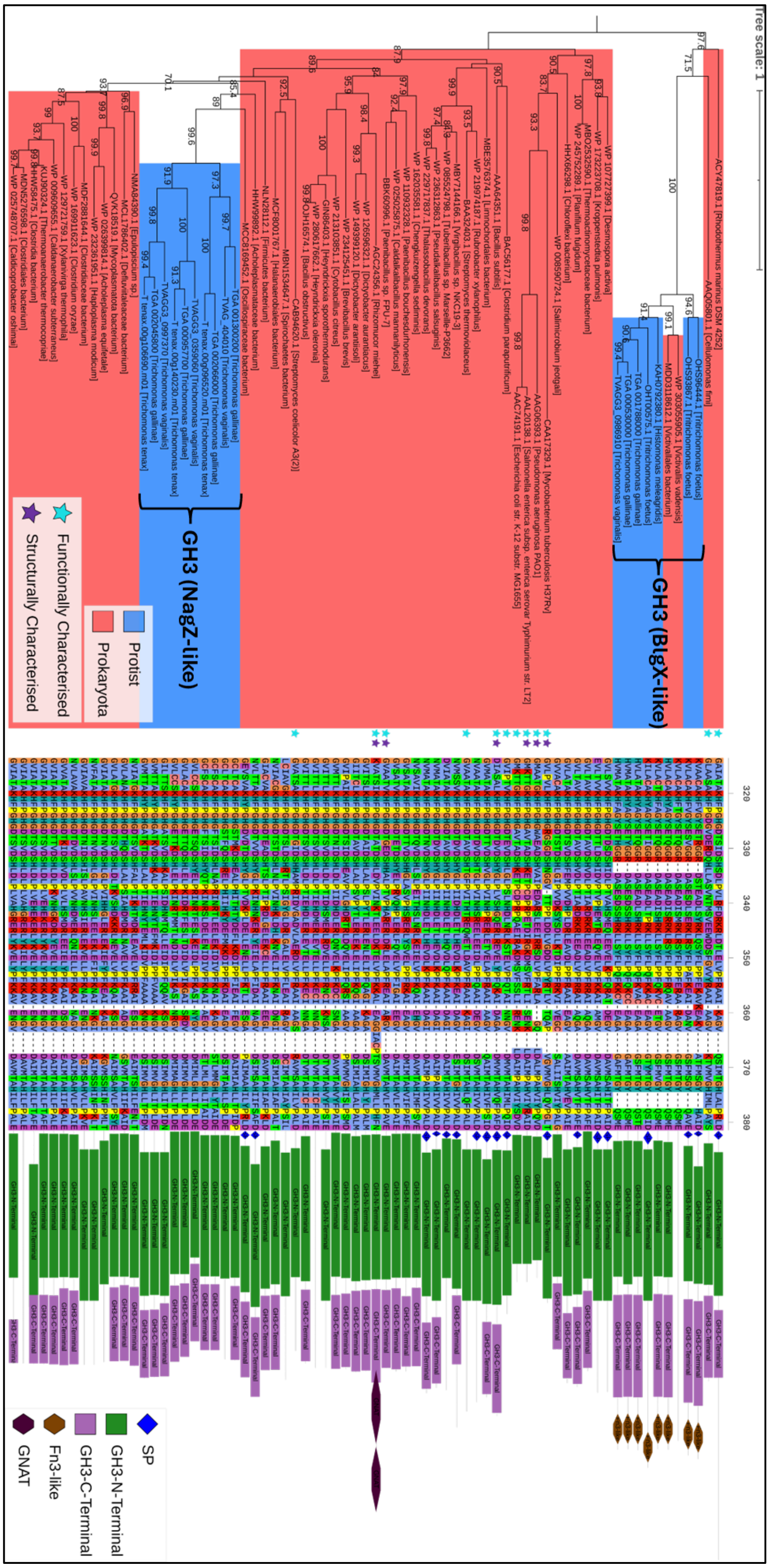
Tree scale: 1
GH3 (BlgX-like)
GH3 (NagZ-like)
Protist
Prokaryota
Functionally Characterised
Structurally Characterised
SP
GH3-N-Terminal
GH3-C-Terminal
Fn3-like
GNAT

**Figure S10: Phylogenetic analysis, multiple sequence alignment and domain architecture of GH3 proteins.** An outgroup-rooted phylogenetic tree was constructed from a masked multiple sequence alignment of Trichomonas and trichomonad GH3 proteins, homologues identified via BLASTp searches against NCBI's nr database and characterized GH3s from the CAZy database. The maximum likelihood phylogeny was inferred using the LG+I+G4 model with ultrafast bootstrapping; bootstrap values >70 are displayed at the corresponding nodes. The scale bar shows the inferred number of substitutions per site. Branch labels include gene ID and species of origin. *Trichomonas* NagZ-like and BlgX-like GH3s are indicated to the right of relevant branches. Clades are colour-coded by taxonomic group: protists (blue) and prokaryotes (red). Functionally characterised proteins are marked with blue stars; structurally characterised proteins with purple stars. A central panel shows a partial multiple sequence alignment, highlighting the conserved NagZ motif. Domain architectures for each sequence are shown to the right of the tree, with domains annotated according to the key in the bottom right of the figure.

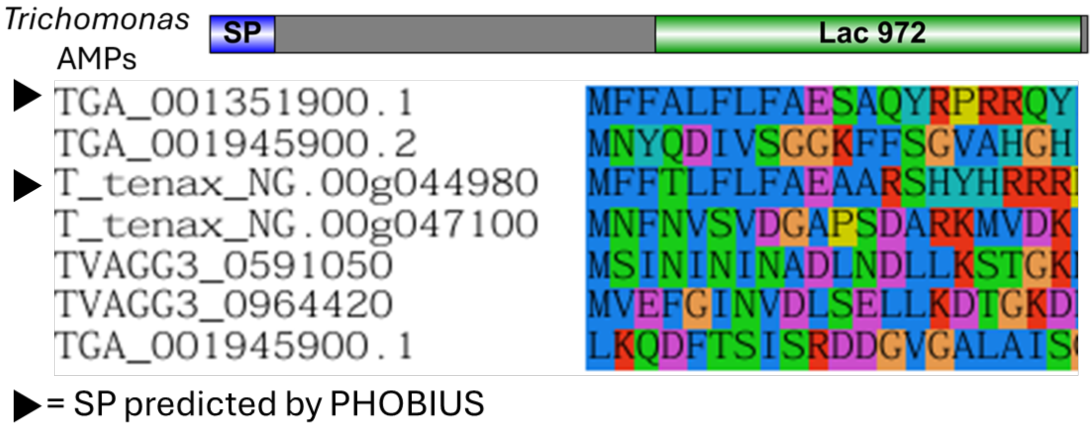


**Figure S11: Domain architecture and N-terminal alignment of *Trichomonas* AMP-Lcn972s.** Top: Domain organisation of *Trichomonas* AMP-Lcn972, with predicted signal peptide (SP) shown in blue and the Lac972 domain in green. Bottom: N-terminal alignment of *Trichomonas* AMP-Lcn972

proteins showing conserved sequence features. Black arrows indicate sequences with PHOBIOUS predicted signal peptides (SP).

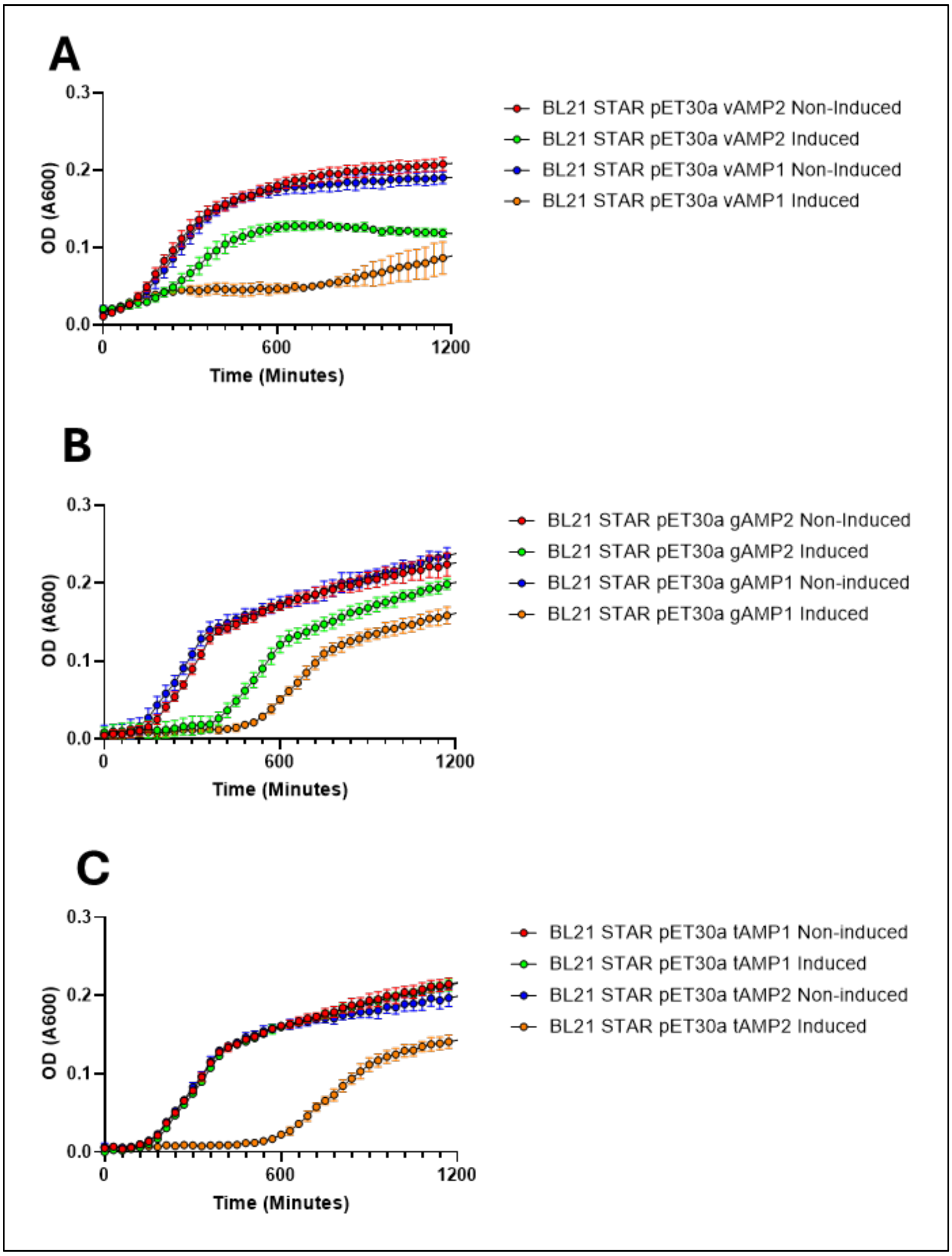


**Figure S12: Induced and non-induced growth curves for BL21 STAR transformed with *Trichomonas* AMP-Lcn972 expression constructs.** Growth curves displaying the mean optical density (A600) of *E. coli* BL21 STAR cultured in LB medium, with or without IPTG induction, while harbouring expression vectors containing *Trichomonas* AMP-Lcn972 inserts from (A) *T. vaginalis*, (B) *T. gallinae*, and (C) *T. tenax*. IPTG was used to induce protein expression. Error bars represent standard deviation. A legend is provided to the right of each panel. N = 8.

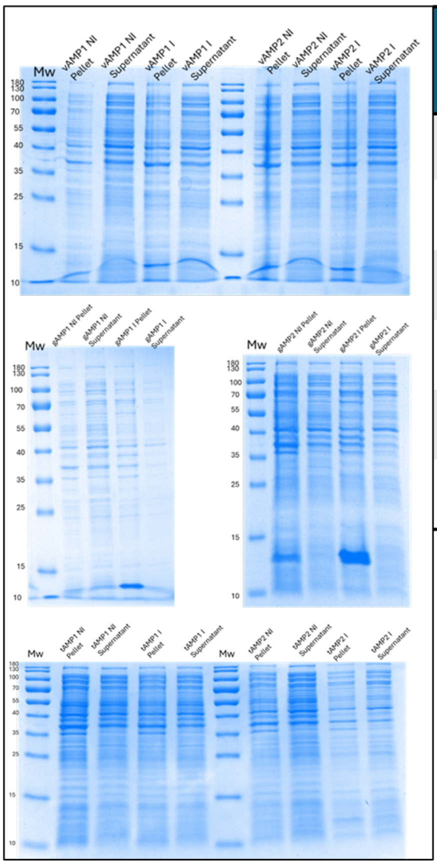


| AMP | Truncated Length in pET30a (AA) | Size with His-tag (kDa) | Constitutively expressed | Expressed with 0.5mM IPTG induction? | Soluble? |
|---|---|---|---|---|---|
| vAMP1 | 78 | ≈13.558 | Yes | Yes | Weakly |
| vAMP2 | 75 | ≈13.051 | Yes | Yes | Weakly |
| gAMP1 | 78 | ≈13.350 | Yes | Yes | Weakly |
| gAMP2 | 78 | ≈13.837 | Yes | Yes | No |
| tAMP1 | 76 | ≈13.214 | No | No | N/A |
| tAMP2 | 78 | ≈13.676 | No | No | N/A |

**Figure S13: SDS-PAGE summary of *Trichomonas* AMP-Lcn972 expression in BL21 STAR *E. coli.*** Left: SDS-PAGE gels displaying protein content from pellet and supernatant fractions following lysis of IPTG-induced (I) and non-induced (NI) *E. coli* BL21 STAR cultures expressing AMP-Lcn972 proteins from *T. vaginalis*, *T. gallinae*, and *T. tenax*. Lane contents are labelled above each respective lane. Right: Summary table outlining the outcomes of the AMP-Lcn972 expression protocol in *E. coli* BL21 STAR, including solubility and expression observations for each construct.

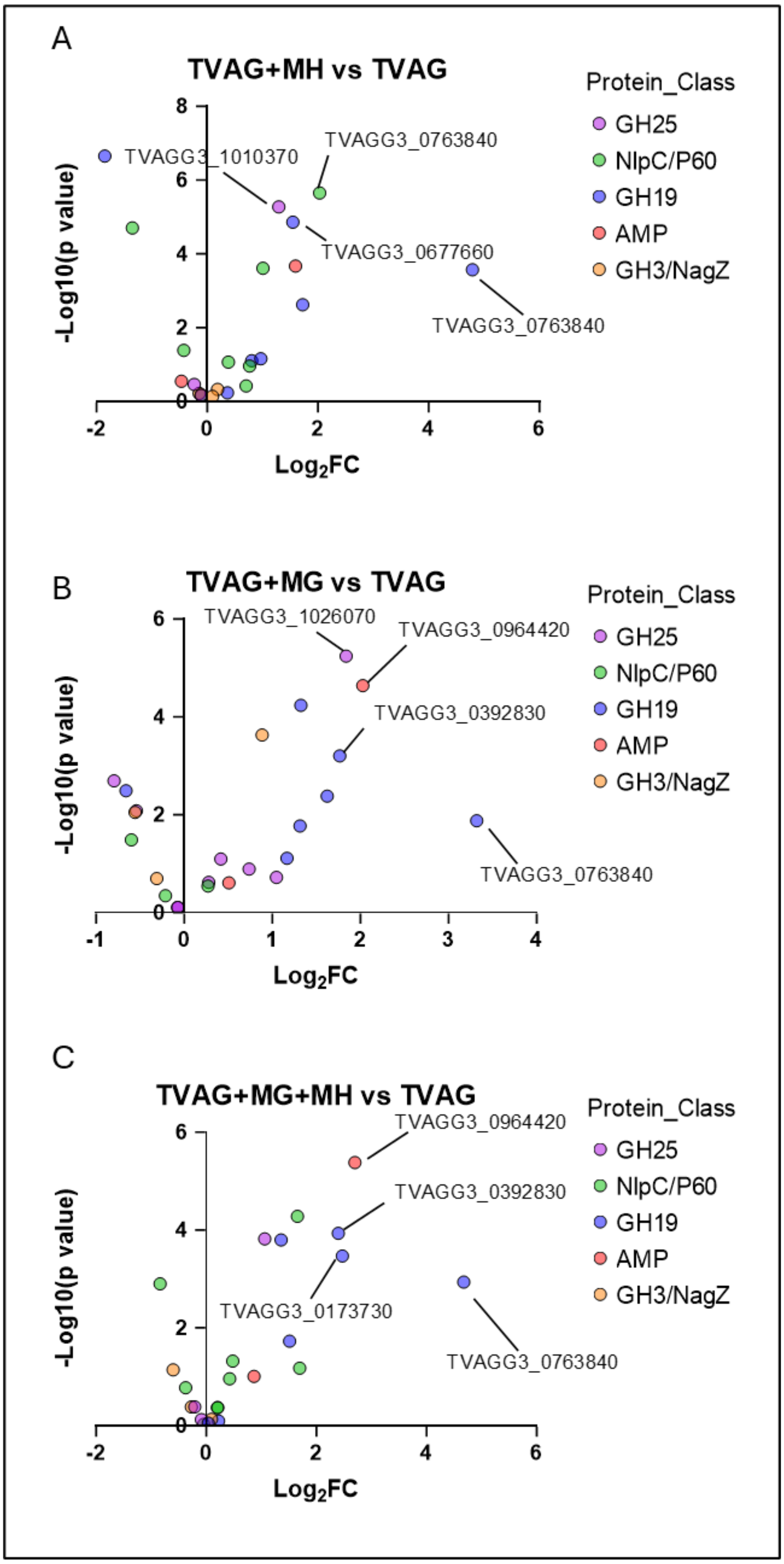


**Figure S14: Differential expression of *T. vaginalis* antibacterial toolkit genes in the presence of *m*ycoplasma-like endosymbionts.** Volcano plots showing differential gene expression ($log_2$ fold change) in *T. vaginalis* when harbouring: (A) a *M. hominis* (MH) endosymbiont, (B) a ca. *M.girerdii*

(MG) endosymbiont, and (C) both endosymbionts concurrently (MG+MH). Points are colour-coded based on the antibacterial toolkit category each gene belongs to, as indicated in the key to the right. Gene IDs are annotated for a selection of the most highly upregulated genes. These RNAseq data used to generate these graphs are from Margarita et al., (2022), see main text.

| BioSample Accession | Sample | Raw reads no. | Raw data (GB) |
|---|---|---|---|
| SAMN37478063 | CC1 | 151,255,726 | 22.7 |
| SAMN37478064 | CC2 | 199,243,678 | 29.9 |
| SAMN37478065 | CC3 | 157,747,480 | 23.7 |
| SAMN37478066 | Tg1 | 215,164,792 | 32.3 |
| SAMN37478067 | Tg2 | 233,888,678 | 35.1 |
| SAMN374780638 | Tg3 | 154,095,748 | 23.1 |
| | Total | 1,111,396,102 | |
| | Average | 185,232,684 | |

**Table S1: Summary of RNAseq reads obtained from *T. gallinae* and *T. gallinae–E. coli* co-culture experiments.** Co-culture samples are labelled with the prefix CC, and monoculture samples with the prefix Tg, each followed by the corresponding replicate number.

| Subject Species Name | Taxonomy (phylum) | BitScore | Query Cover | E value | % identity | Accession |
|---|---|---|---|---|---|---|
| *Tritrichomonas foetus* | Parabasalia | 89.7 | 78% | 3.00E-18 | 41.67 | XP_068363938.1 |
| *Trichomonas vaginalis G3* | Parabasalia | 80.9 | 75% | 2.00E-16 | 41.75 | XP_001329491.1 |
| *Trichomonas vaginalis G3* | Parabasalia | 79.7 | 73% | 5.00E-16 | 44.12 | XP_001325452.1 |
| *Histomonas meleagridis* | Parabasalia | 71.2 | 70% | 3.00E-12 | 36.46 | KAH0788655.1 |
| *Histomonas meleagridis* | Parabasalia | 70.9 | 70% | 3.00E-12 | 36.46 | XP_067770210.1 |
| *Streblomastix strix* | Parabasalia | 55.8 | 74% | 9.00E-07 | 30.39 | KAA6363257.1 |
| *Cellulomonas sp. KRMCY2* | Actinomycetota | 47.4 | 68% | 0.001 | 34.04 | WP_051480476.1 |
| *Kocuria sp. 36* | Actinomycetota | 46.2 | 50% | 0.004 | 39.13 | WP_129659434.1 |
| *Monocercomonoides exilis* | Preaxostyla | 46.2 | 73% | 0.005 | 26.73 | XP_067727518.1 |
| *Cellulomonadaceae bacterium* | Actinomycetota | 45.1 | 54% | 0.011 | 31.08 | NTW40279.1 |
| *Streptomyces sp. NPDC023588* | Actinomycetota | 45.1 | 48% | 0.011 | 39.39 | MEU4731520.1 |

**Table S2: BLASTp hit list for *Trichomonas* AMP-Lcn972 protein TGA_001351900.1 against the NCBI nr database.** The table lists the top hit per species, including associated accession numbers, taxonomic classification, BitScore, query coverage, E-value, and percentage identity.

| Gene | Forward Primer (5'-3') | Reverse Primer (5'-3') |
|---|---|---|
| TGA_001274700.1 | CGCGCGGCAGCCATATTGGCCATAGAATGCCAGAATC | GGTGGTGGTGCTCGATTAGAAAATACCGCAAGCTCTG |
| TVAGG3_0677660 | CGCGCGGCAGCCATAACATGCAACCTGAATCCAACG | GGTGGTGGTGCTCGAATGCTGAAGGATTTAAGAGGG |
| TGA_001274700.1 MUT1 [E202Q] | GCCGCAGGCTGATTGGTGAGAGCATTGAG | CTCAATGCTCTCACCAATCAGCCTGCGGC |

**Table S3: PCR and mutagenesis primers for *Trichomonas* GH19 genes.** Primers used for PCR amplification of *T. gallinae* GH19 (TGA_001274700.1), *T. vaginalis* GH19 (TVAGG3_0677660), and site-directed mutagenesis of *T. gallinae* GH19 (TGA_001274700.1 MUT1 [E202Q]).
